# Neuronal compartmentalization results in “impoverished” axonal mitochondria

**DOI:** 10.1101/2025.10.27.684882

**Authors:** Adrian Marti Pastor, Liam S.C. Lewis, Stephan Müller, Pedro Guedes-Dias, Lorena Olifiers, Polina Pelkonen, Almudena del Rio Martin, Caroline Fecher, Nikolai P. Skiba, Ying Hao, Antoneta Gavoci, Laura Trovo, Mehmet Akif Aktas, Hariharan M. Mahadevan, Angelika B. Harbauer, Howard M. Bomze, Vadim Y. Arshavsky, Monika S. Brill, Romain Cartoni, Stefan F. Lichtenthaler, Sidney M. Gospe, Thomas Misgeld

**Affiliations:** Institute of Neuronal Cell Biology, School of Medicine and Health, Technical University of Munich (TUM), Munich, Germany; German Centre for Neurodegenerative Diseases (DZNE), Munich, Germany; Department of Pharmacology and Cancer Biology, Duke University School of Medicine, USA; Neuroproteomics, School of Medicine and Health, TUM University Hospital, TUM, Munich, Germany; Department of Ophthalmology, Duke University School of Medicine, USA; Max Planck Institute for Biological Intelligence, Martinsried, Germany; Munich Cluster for Systems Neurology, Munich, Germany

**Author notes:** Correspondence to: T.M. Department of Cell Biology & Physiology, Washington University School of Medicine, St. Louis, MO, USA. These authors contributed equally (Adrian Marti Pastor, Liam S.C. Lewis).

## Abstract

Mitochondria differ depending on their location within a neuron. Morphological heterogeneity between somatic, dendritic, and axonal mitochondria is well established. Emerging evidence suggests that further specialization is needed to meet the unique demands of different neuronal compartments. However, the molecular and functional diversity of mitochondria within a neuron remains poorly understood. Here, we utilized proteomics in MitoTag mice to profile somatodendritic and axonal mitochondria across four distinct neuron types, thereby generating a compendium of intracellular mitochondrial diversity. Combining proteomics, functional, and immunofluorescence analyses, we demonstrated that axonal mitochondria are not defined by the presence of unique proteins, but rather by the selective loss or preservation of specific pathways compared to their somatodendritic counterparts. This results in “impoverished” axonal mitochondria, which are characterized by diminished mtDNA expression and impaired oxidative phosphorylation yet retain other pathways, such as fatty acid metabolism. Bioinformatic analyses of multiomic data identified local translation as one mechanism underlying compartment-specific diversity. Together, these findings provide a comprehensive *in vivo* framework for understanding mitochondrial specialization across neuronal compartments.

## Introduction

Mitochondria play many roles in cells, including energy provision, metabolic regulation, calcium buffering, and apoptosis (Misgeld & Schwarz, 2017). Such multifaceted functions can sometimes be competitive, e.g., with regard to bioenergetic vs. biosynthetic roles (Ryu et al., 2024). Consequently, a single mitochondrion may be unable to fulfill all required roles simultaneously. A solution to this challenge is to diversify mitochondria, for example, via their proteome. A cell would then host diverse mitochondrial subpopulations with an adapted proteome to meet different functional demands. Evidence of intracellular mitochondrial diversity has been shown, for example, in adipocytes (Benador et al., 2018), myocytes (Kuznetsov et al., 1998), and pancreatic acinar cells (Park et al., 2001).

In neurons – the highly polarized cells of the nervous system – such mitochondrial diversity seems essential: Different neuronal compartments, such as dendrites, soma, and axon, have diverse bioenergetic and biochemical needs (Pekkurnaz & Wang, 2022). For instance, postsynaptic processes have especially high energy demands, while at the presynapse, biosynthetic processes, e.g. related to neurotransmitter synthesis, might be more relevant (Harris et al., 2012) Moreover, it is unclear whether a complete mitochondrial proteome can be maintained in an axon. Many axonal mitochondria reside a day’s transport journey away from the nucleus, where more than 99% of mitochondrial proteins are encoded (Misgeld & Schwarz, 2017). While some mitochondrial proteins are exceptionally long-lived, others would not “survive” this journey (Bomba-Warczak et al., 2021). A subset of these short-lived proteins is maintained by a dedicated local translation system (Harbauer et al., 2022). However, whether this suffices to endow mitochondria in all parts of a neuron with the same proteome composition remains to be shown. Deviation from a “complete” mitochondrial proteome in the axon, either due to functional demands or homeostatic constraints, may have important implications for compartmental vulnerability in axonopathic diseases, such as optic neuropathy, Charcot-Marie-Tooth disease, amyotrophic lateral sclerosis, or hereditary spastic paraplegia (Coleman, 2005).

Indeed, mitochondria in different compartments of the neuron, e.g., its dendrites versus the axon, show morphological and ultrastructural heterogeneity (e.g., Faitg et al., 2021; Mendelsohn et al., 2022). The functional implications of these unique shapes are debated. Several lines of argument, however, suggest that mitochondria play differential roles in axons or synapses, related to their efficiency in energy provision, substrate utilization, and calcium buffering capacity (Akefe et al., 2024; Hirabayashi et al., 2024; Jang et al., 2025). However, it remains to be proven whether such specialized axonal functionality is the result of local molecular diversification or rather a consequence of the mitochondrion’s environment (local redox equilibria, metabolite availability, or ionic driving forces), which imposes non-canonical functionality on a canonical molecular machinery.

Neurons present a unique opportunity to study intracellular mitochondrial diversity, as their axons can span considerable distances away from the soma and dendrites, allowing physical separation. However, the mechanisms by which neuronal mitochondria diversify remain poorly understood, mainly due to three challenges: (1) the difficulty of fully differentiating neurons *in vitro*, which is known to impact axonal mitochondrial function (e.g. with regards to dynamics; Faits et al., 2016; Misgeld & Schwarz, 2017; Smit-Rigter et al., 2016) and limits the use of otherwise powerful approaches to neuronal compartmentalization, such as microfluidic chambers (Taylor et al., 2010), (2) the lack of tools that allows for the selective isolation of neuronal mitochondria – which contrasts with other organelles, where tools such as the RiboTag have revealed unique local functionality (in this case related to diversifying the compartmental translatome of neurons; Shigeoka et al., 2016); (3) the problem to clearly phenotype mitochondrial pools *in situ*, since for example metabolic biosensors cannot easily distinguish between organelle diversification versus environmentally imposed differences. To circumnavigate these challenges, previous attempts to understand mitochondrial specialization used isolation and proteomic analysis of synaptosomes (e.g., Graham et al., 2017; Van Oostrum et al., 2023). However, in such studies, a clean comparison with non-synaptic neuronal mitochondria is difficult. The non-synaptic “background” proteome of the whole tissue, from which the synaptosomes are isolated and to which they are referenced, invariably contains organelles from several cell types. Therefore, intracellular diversity between synaptic and other neuronal mitochondria is convolved with and obscured by the substantial intercellular mitochondrial diversity, e.g., between neurons and glia (Fecher et al., 2019). Complementary approaches that would resolve this problem by cell-type-specific protein tagging have not yet been used for this purpose, in part because even for whole cell proteomes, they have shown limited quantitative or qualitative coverage of mitochondrial proteins (Alvarez-Castelao et al., 2017; Hobson et al., 2022). To our knowledge, a systematic approach comparing mitochondrial populations from subcellular compartments of a single neuron type therefore remains elusive.

In this study, we introduce a compartmentalized MitoTag-proteomics approach that overcomes these problems and leverages neuroanatomy to reveal subcellular mitochondrial diversity across four neuronal cell populations. Our study provides insights into the mitochondrial specializations in the somatodendritic versus the axonal compartment. This revealed axonal mitochondria as an “impoverished” version of their somatodendritic counterparts, meaning that these distal organelles can only maintain a part of a “complete” mitochondrion’s proteome and functionality.

## Results

### Somatodendritic and axonal mitochondria can be separated in MitoTag mice

We adapted the MitoTag approach (Fecher et al., 2019) to profile mitochondria from different neuronal compartments *ex vivo*, specifically the somatodendritic and axonal parts of nerve cells (Fig. 1A). For this, we chose four genetically targetable neuron types in which the axonal projections are well separated from the somatodendritic region. We crossed the corresponding Cre-driver lines to the *floxed* MitoTag reporter strain, to express GFP on the outer mitochondrial membrane (GFP-OMM; see Methods): (1) layer V cortical projection neurons (CPN), labelled using Rbp4-Cre, have their somata and dendrites in the cortex and project axons through the striatum (Fig. 1B); (2) central cholinergic neurons (CCN; ChAT-Cre), reside in basal forebrain nuclei and striatum, and send axons to the cortex (Fig. 1C); (3) motor neurons (MN; ChAT-Cre), are located in the spinal cord and project long axons through the sciatic nerve (Fig. 1D); and (4) retinal ganglion cells (RGC; VGlut2-Cre), have somata and dendrites in the retina and project axons through the optic nerve (Fig. 1E). We confirmed that these tissues are enriched in compartment-specific mitochondria with high-resolution imaging (Fig. 1F-M).

**Fig. 1:**
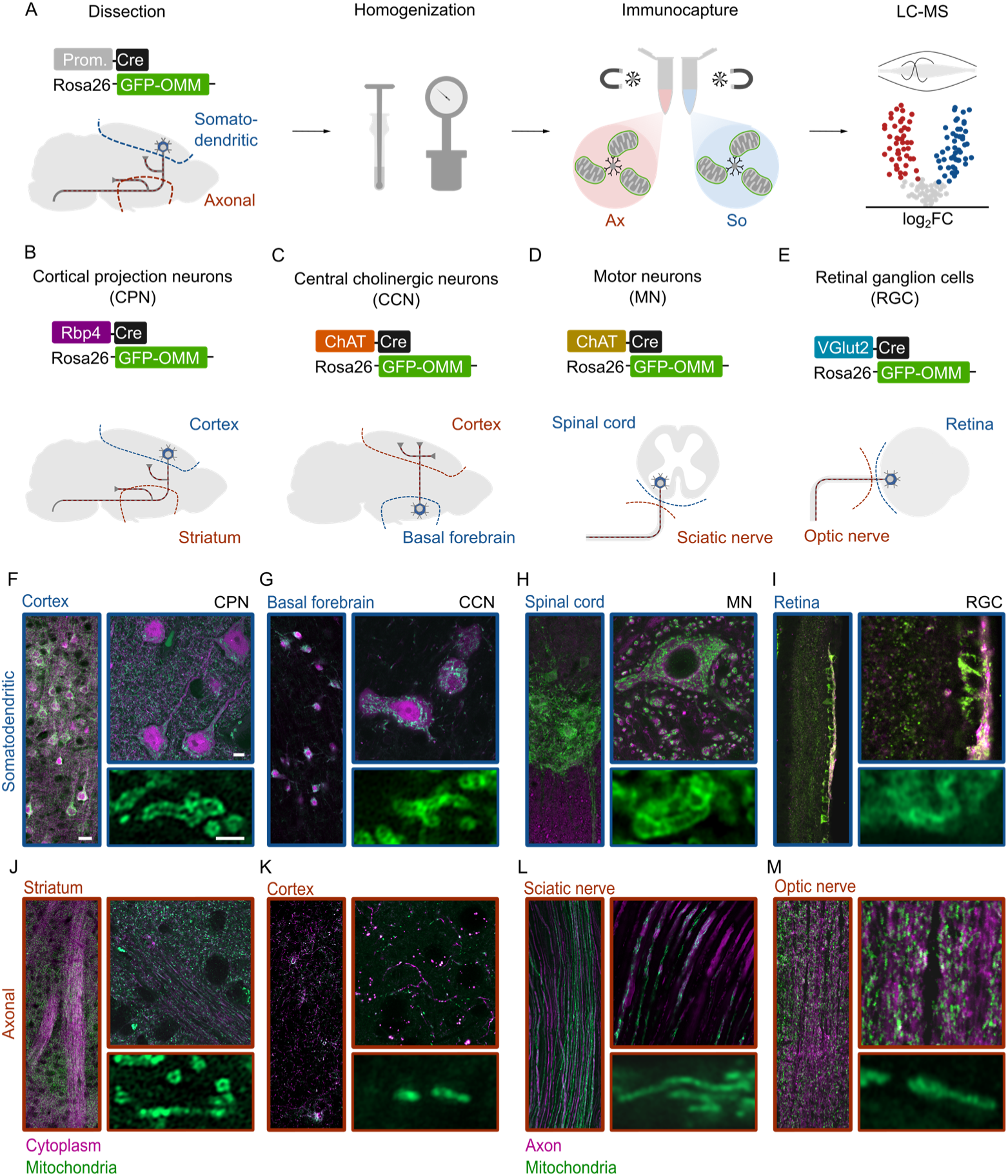
Compartmentalized mitochondria in neuronal MitoTag mouse lines. **A**. Schematic of the immunocapture of cell-type-and compartment-specific mitochondria using MitoTag. From left: dissection of the brain regions of interest, mechanic and pressure-based homogenization, immunocapture of compartment-specific mitochondria with magnetic beads, liquid chromatography-mass spectrometry followed by data analysis. **B-E**. Schematic of the dissection strategy and transgenic mouse lines to obtain mitochondria from either the somatodendritic or axonal compartment from four neuronal populations: (**B**) Layer V cortical projection neurons (CPN) labeled with Rbp4-Cre/GFP-OMM, (**C**) Central cholinergic neurons (CCN) labeled with ChAT-Cre/GFP-OMM, (**D**) Motor neurons (MN) labeled with ChAT-Cre/GFP-OMM, and (**E**) Retinal ganglion cells (RGC) labeled with VGlut2-Cre/GFP-OMM. **F-M**. Airyscan and confocal images of the tissues selected for dissection and immunocapture. GFP-OMM labelled mitochondria displayed in green and cell-type specific cytoplasm labelled with Cre-dependent tdTomato expression (CPN and CCN; magenta) or tissue axons labelled with tubulin-ß3 immunostaining (MN and RGC; magenta). (**F-I**) somatodendritic or (**J-M**) axonal compartment. Scale bars: Left overview panels, 25 µm; right details panels, 5 µm (upper); 1 µm (lower).

We decided to use several systems in parallel for several reasons: (1) We aimed to identify a “consensus” set of proteomic differences between mitochondrial populations across a variety of neuron classes, e.g. ones that have axonal projections in the CNS versus PNS (CPN, CCN and RGC vs. MN), are myelinated versus non-myelinated (CPN, MN, RGC vs. CCN), and ones that carry presynaptic terminals in the dissected regions versus ones that do not (CPN, CCN vs. MN, RGC). (2) We also wanted to cancel out the unique technical challenges that each system presents, including variable tissue sizes and densities of labeled mitochondria, as well as different sources of potential contamination. While in all systems the axon initial segment and recurrent collaterals are comprised in the somatodendritic isolate, e.g. in cortex and retina, additionally small populations of cholinergic interneurons exist (von Engelhardt et al., 2007; L. Wang et al., 2020), the mitochondria of which will be captured as axonal; similarly, a small subset of cholinergic sensory projections exist in the sciatic nerve, resulting in a contribution of a “non-classical” pseudo-unipolar projections to the *bona fide* motor axons (Tata et al., 2004). (3) Finally, our approach also resolves the problem that immunocapture (IC) can be susceptible to background effects, meaning the influence of the tissue from which an IC isolate originates. The inverse symmetry of CPN and CCN neurons allows us to cancel out this confounder via a cross-over design, where the somatic background tissue of one neuron type is the axonal background tissue of the other, and vice versa.

First, we ensured that axonal mitochondria can be effectively immunocaptured, as they are smaller and more compact than their somatodendritic counterparts, potentially exposing less of the target antigen (Faitg et al., 2021; Fischer et al., 2018; Lewis et al., 2018). We focused on the axonal compartment of RGCs, which we deemed the most challenging system due to its small tissue size in mice. Electron micrographs confirmed the successful immunocapture of axonal RGC mitochondria, which retained their characteristic smaller size relative to those in the somatodendritic compartments (Supp. Fig. 1A-B). Isolated axonal mitochondria expressed endogenous mitochondrial proteins as well as the GFP-outer mitochondrial membrane (OMM) tag (Supp. Fig. 1C). To control for background contamination, we mixed optic nerves from a GFP-OMM-expressing mouse, which in addition carried a marker mutation (Ndufs4 knockout), with those from a wild-type mouse. Only mitochondria from the knockout mouse were recovered, indicating high capture specificity (Supp. Fig. 1D-E).

### Somatodendritic and axonal mitochondria exhibit a specialized proteome

We compared the differential expression of mitochondrial proteins between somatodendritic and axonal compartments across these neuronal populations using mass spectrometry (Fig. 2A). The resulting compartment-specific mitochondrial proteomes (immunocapture using anti-GFP; IC-GFP) achieved high coverage of the MitoCarta3.0 compendium (Supp. Fig. 2A) of neurons in the brain (average of IC-GFP CPN and CCN isolates: 77.6% ± 0.3; mean ± SEM). This was in a similar range to the mitochondrial proteome coverage obtained using a pan-mitochondrial isolation (targeting Tom22; IC-TOM; 81.6% ± 0.2). Coverage decreased with tissue size (spinal cord: 59.9% ± 4.7; sciatic nerve: 28.0% ± 5.1; retina: 14.8% ± 1.4; and optic nerve: 13.3% ± 0.7). The sub-mitochondrial localization of quantified proteins was uniformly distributed across samples and compartments (Supp. Fig. 2B). Additionally, when analyzed collectively, principal component analysis (PCA) revealed three major clusters from proteomes: from brain, RGC and MN samples, possibly reflecting intrinsic tissue similarities or batch effects (Supp. Fig. 2C); however, clustering of individual datasets revealed clear compartmental separation (Supp. Fig. 2D-G). Lastly, we assessed the enrichment quality of our immunocapture by probing neuronal and astrocytic mitochondrial markers (Fecher et al., 2019) in neuron-derived mitochondria (IC-GFP) relative to mitochondria from their background tissue (IC-TOM; Supp. Fig. 2H). As expected, mitochondria isolated from CPN neurons showed strong enrichment of neuron-specific mitochondrial proteins and depletion of astrocytic marker proteins, indicating high specificity for neuronal mitochondria. CCN mitochondria exhibited less distinct enrichment, suggesting lower immunocapture efficiency in line with the overall smaller fraction of labelled and captured mitochondria in this sample (Fecher et al., 2019).

**Fig. 2:**
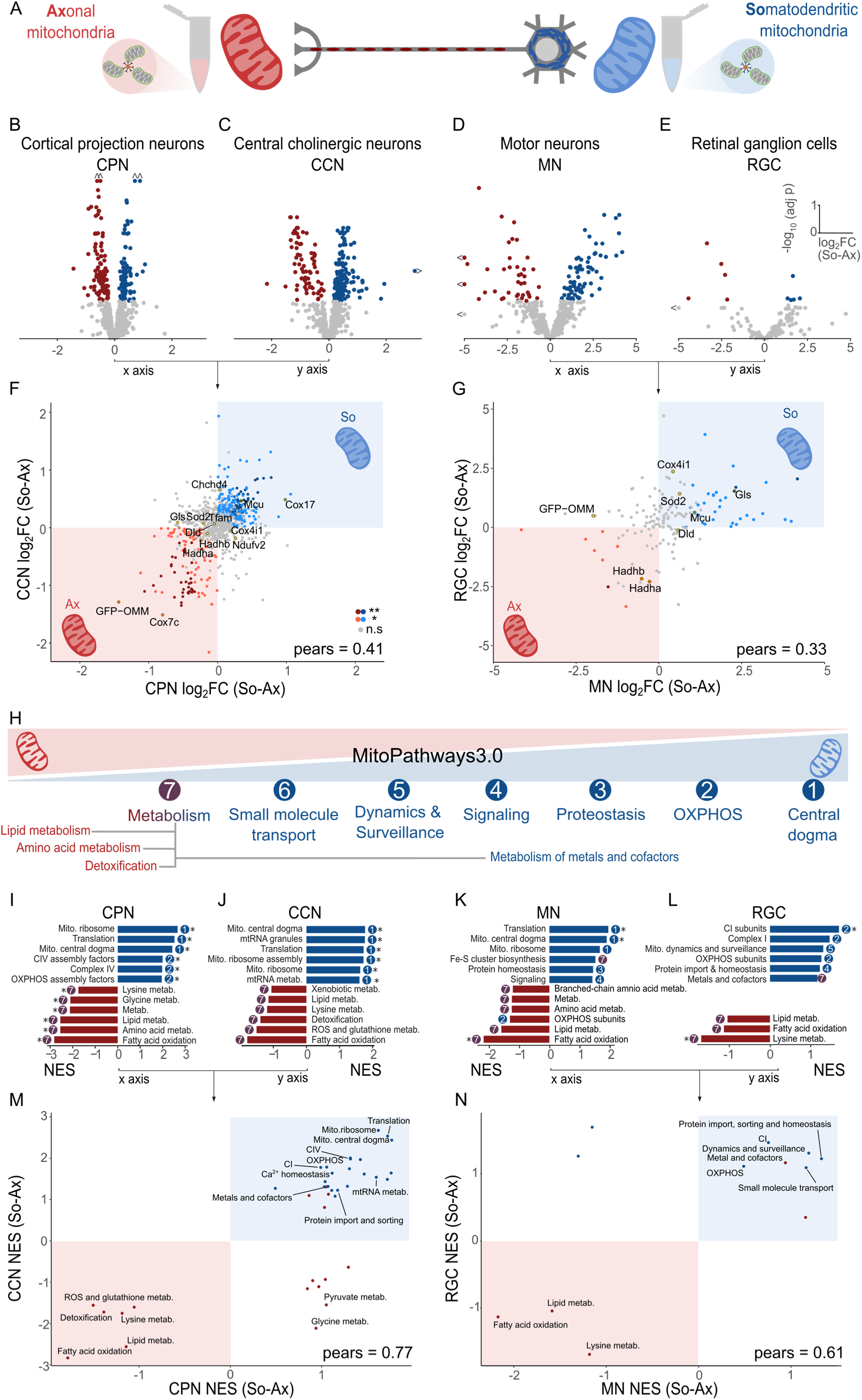
Proteomic specialization of compartment-specific mitochondria. **A.** Axonal and somatodendritic mitochondria are isolated from the same neuronal population. **B-E**. Differential expression analysis (volcano plots) comparing cell-type- and compartment-specific mitochondrial proteomes between compartments (So/Ax) for (B) CPN, n = 17, (C) CCN, n = 13, (D) MN, n = 6, (E) RGC, n=6. Tested with moderated t-test (limma model); adjusted p-value (adj p, FDR) < 0.05 for significance; displayed as soma-enriched (blue), axon-enriched (red), or unchanged (non-significant; gray). **F-G**. Correlation scatterplots of proteins merging the log_2_ fold-changes (log_2_FC) displayed in plots B-E; (F) x-axis: So/Ax from CPN, y-axis: So/Ax from CCN, and (G) x-axis: So/Ax from MN, y-axis: So/Ax from RGC. Colored as differentially localized in both neurons (**: somatodendritic, dark blue; axonal, dark red), only one of them (*: somatodendritic, light blue; axonal: light red), or unchanged (n.s.: discordant or non-significant, grey). **H.** Summarizing scheme of the relative enrichment of compartmentalized mitochondrial pathway groups shared across neuron types as per panels I-N. **I-L**. Gene set enrichment analysis (GSEA) with MitoPathways3.0: Somatodendritically-(blue) and axonally-enriched (red) pathways labelled with the pathway group they belong to from panel H and * for significance: FDR < 0.05. **M-N**. Correlation scatterplots of pathway normalized enrichment score (NES) from panels I-L; (M) x-axis: So/Ax from CPN, y-axis: So/Ax from CCN, and (N) x-axis: So/Ax from MN, y-axis: So/Ax from RGC. Pears: Pearson correlation coefficient; So: somatodendritic, Ax: axonal, FC: fold change, CI: complex I, CIV: complex IV, metab.: metabolism, mito.: mitochondrial.

Comparative analysis of mitochondrial proteins across compartments revealed distinct proteomes, with 110 and 114 differentially enriched proteins in axonal and somatodendritic mitochondria from CPN, respectively (Fig. 2B), 73 and 182 in CCN (Fig. 2C), 41 and 80 in MN (Fig. 2D), and 5 in RGC for both compartments (Fig. 2E). Despite possible cell-type-specific differences, there was substantial overlap in the classification of proteins as somatodendritic or axonal, and enrichment ratios were correlated across cell types, indicating generalizability of the proteomes, as shown by correlation coefficient factors of Pearson r = 0.41 (p < 0.001) for the brain data (Fig. 2F) and Pearson r = 0.33 (p < 0.001) for the nerve data (Fig. 2G). Compartmentalized mitochondria also showed a striking similarity at the pathway level (Fig. 2H-N): somatodendritic mitochondria were enriched in proteins involved in mtDNA maintenance, transcription, and translation (i.e., mitochondrial central dogma), oxidative phosphorylation, metal and cofactor metabolism, and proteostasis. In contrast, axonal mitochondria were enriched in proteins related to fatty acid oxidation (FAO) and amino acid metabolism. Pathway-level enrichment ratios substantially improved the correlation coefficients, with Pearson r = 0.77 (p < 0.001) for CPN-CCN and Pearson r = 0.61 (p < 0.05) for MN-RGC correlation (Fig. 2M-N, Supp. Fig. 2I), indicating a general agreement across cell types.

Non-mitochondrial proteins are also identified in our proteomes but their interpretation requires caution, as these proteins may represent true intracellular interactors of mitochondria or contaminants from the cell lysate. Supporting the former in our most robust proteome, non-mitochondrial proteins from the somatodendritic compartment of the CPN were significantly enriched in microbody, peroxisomal, ribosomal, and spliceosome proteins, with the latter indicating proximity to the nucleus. Non-mitochondrial proteins from the axonal compartment showed a trend toward enrichment in axonal and presynaptic proteins (Supp. Fig. 2J).

To further investigate spatial proteomic dynamics, we collected mitochondrial proteomes from distal CPN axons in the dorsal corticospinal tract of the spinal cord (Supp. Fig. 3A-B). Integrating these with proximal axonal mitochondria from the striatum and somatodendritic mitochondria from the cortex allowed us to track protein abundance changes as a function of distance from the soma (Supp. Fig. 3C). Linear regression modeling revealed protein- and pathway-level results consistent with previous findings. We found 184 proteins depleted along the CPN axon and 378 proteins that increase their levels, with slope values reflecting the rate of protein acquisition or depletion along the axon (Supp. Fig. 3D). Using the slope as the quantitative value, pathways related to mtDNA expression were the main hit for somatodendritic mitochondria, while FAO as well as amino acid metabolism stood out for axonal mitochondria (Supp. Fig. 3E).

### Axonal mitochondria show reduced oxidative phosphorylation but preserve fatty acid oxidation

To functionally validate our proteomic findings, we immunocaptured mitochondria from the somatodendritic and axonal compartments of the CPN, due to the high quality and yield of these mitochondria. We assessed their respiratory activity using both FAO and tricarboxylic acid (TCA) cycle substrates. Overall, respiration was lower in axonal than in somatodendritic mitochondria (Fig. 3A). No significant differences were observed when FAO was coupled to oxidative phosphorylation (0.92 ± 0.26, p = 0.65). By contrast, TCA-driven respiration was lower in axonal than in somatodendritic mitochondria (primarily feeding into Complex II (CII): 0.72 ± 0.08, p = 0.03; or Complex I (CI): 0.83 ± 0.10, p = 0.16, respectively). Furthermore, uncoupled respiration, which reflects maximum respiratory capacity, was significantly diminished in axonal mitochondria (0.76 ± 0.08, p = 0.03) (Fig. 3B), underscoring bioenergetic compartmentalization. When comparing tissue-level mitochondrial populations using IC-TOM, striatal mitochondria (the tissue where CPN axons are located: axonal “background”) displayed non-significant but consistently higher respiration across TCA (1.27 ± 0.31 in CI, p = 0.42; 1.36 ± 0.15 in CII, p = 0.31), FAO (1.26 ± 0.21, p = 0.37), and uncoupled conditions (1.31 ± 0.18, p = 0.31) relative to cortical mitochondria (somatodendritic “background”), further demonstrating that the respiration difference between the CPN compartments are not attributable to “background” mitochondrial populations (Supp. Fig. 4A-B).

**Fig. 3:**
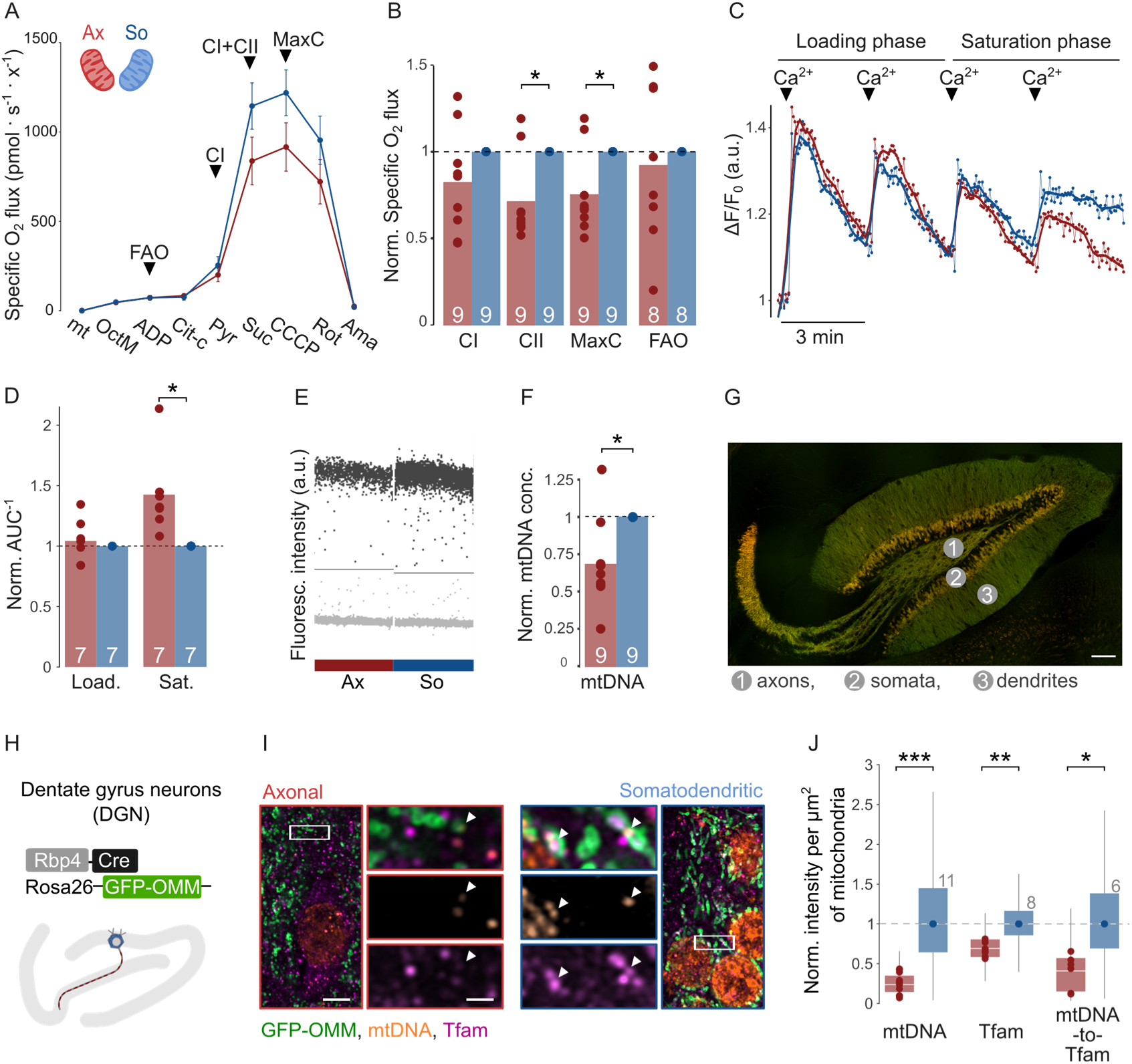
Functional validation of proteomic results and mtDNA quantification. **A**. Respirometry curve of CPN isolated mitochondria from somatodendritic (blue) or axonal compartments (red). Points are averages with standard errors. Arrowheads indicate fold change quantification for panel B. Mt: mitochondria, OctM: octanylcarnitine and malate, ADP: adenine diphosphate, Cit-c: cytochrome c, Pyr: pyruvate, Suc: succinate, CCCP: Carbonyl cyanide m-chlorophenyl hydrazone, Rot: rotenone, Ama: antimycin A. **B**. Pair-wise normalized replicates from panel A. CI: complex I, CII: complex II, MaxC: maximum capacity, FAO: fatty acid oxidation. **C**. Example curve of mitochondrial calcium uptake assay using CPN isolations. Arrowheads indicate the addition of calcium. **D**. Pair-wise normalized replicates from calcium uptake assay shown as AUC^-1^. **E**. Representative quantification of mitochondrial DNA (mtDNA) using digital PCR, separating positive (black) and negative (gray) partitions in somatodendritic (So) and axonal (Ax) CPN mitochondria (horizontal line). **F**. Pair-wise normalized replicates from dPCR. **G**. Airyscan image of the dentate gyrus neurons (DGN) as labeled by Rbp4-Cre driven GFP-OMM (green), and tdTomato (orange). Neuro compartmentalization labelled. Scale bar 100 µm. **H**. Schematic view of mouse transgenic line and DGN in the hippocampus. **I**. Representative Airyscan images of co-localization of mtDNA (orange), Tfam (magenta), and GFP-OMM-tagged mitochondria (green) in the axonal (red boxes) and somatodendritic (blue boxes) areas of the DGN. Arrowheads point to mtDNA nucleoids inside mitochondria with or without Tfam co-localization. Scale bar: 5 µm (left), 1 µm (right). **J**. Quantification of immunofluorescence intensity of mtDNA, Tfam, and their ratio inside DGN mitochondria. Boxes show overall instensity distribution in mitochondrial area, the points show medians per mouse. *: p < 0.05, **: p < 0.01, ***: p < 0.001. Blue: somatodendritic (So) mitochondria, red: axonal (Ax) mitochondria.

### Neuronal compartments differ in mitochondrial calcium buffering capacity

To further assess the functional implications of our proteomic data, we examined the calcium uptake capacity of isolated compartment-specific mitochondria from CPN. In our pathway analysis, calcium handling was modestly enriched in somatodendritic mitochondria (Fig. 2H, M. Signaling). Indeed, somatodendritic mitochondria displayed higher protein abundance for most mitochondrial calcium uniporter complex (MCUC) subunits (Supp. Fig. 4C). To assay calcium buffering, we resuspended the mitochondrial isolates in a calcium indicator-containing medium and monitored the uptake fluorometrically after the addition of calcium. The assay was divided into two distinct phases: a loading phase (first two calcium additions) and a saturation phase (third and fourth additions; Fig. 3C). During the loading phase, axonal mitochondria exhibited identical calcium uptake capacities to somatodendritic mitochondria (1.04 ± 0.07, p = 0.55). Upon calcium overload in the saturation phase, however, axonal mitochondria buffered significantly better than their somatodendritic counterparts (1.43 ± 0.13, p = 0.02) (Fig. 3D). To potentially explain this discrepancy, we explored the stoichiometric of the MCUC in the two compartments. While most MCUC components had the same expression levels relative to Mcu, Emre – an essential determinant of MCU complex stability and function (Sancak et al., 2013; Y. Wang et al., 2019) – was relatively over-abundant in axonal mitochondria (Supp. Fig. 4C). This could stabilize the MCUC, allowing superior calcium uptake performance of axonal mitochondria during the saturation phase. When comparing striatal and cortical regional mitochondrial populations (axonal and somatodendritic background, respectively), striatum mitochondria exhibited higher calcium uptake capacity (1.45 ± 0.49, p = 0.43, normalized to cortical mitochondria). The uptake was Ruthenium 360-sensitive, confirming MCUC dependency (Supp. Fig. 4D-E).

### Axonal mitochondria contain less mitochondrial DNA

To assess mitochondrial DNA (mtDNA) levels in different neuronal compartments, we used digital PCR (dPCR) to quantify mtDNA amplified from mitochondria isolated from the somatodendritic and axonal regions of CPNs. We targeted the mitochondrial NADH-ubiquinone oxidoreductase chain 1 (mt-Nd1) gene locus and validated the specificity of amplification using the Cytochrome b (Cytb) gene as a second mtDNA locus. Indeed, this analysis revealed a significant reduction (0.69 ± 0.1, p = 0.01) of mtDNA in axonal mitochondria, confirming our pathway analysis results (Fig. 3E-F). Additionally, average striatal mitochondria (axonal “background”) contained slightly higher (1.18 ± 0.26, p = 0.52) mtDNA copy numbers than cortical mitochondria (somatodendritic “background”; Supp. Fig. 4F-G). Importantly, we found a strong correlation in mtDNA quantification with our two primer pairs (mt-Nd1 and Cytb loci; Pearson r = 0.999, p < 0.001), indicating that the mtDNA was intact and amplification was robust (Supp. Fig. 4H). Additionally, our mitochondrial isolation protocol achieved a high enrichment (1628 ± 122-fold) of mtDNA over nuclear DNA (nDNA; probed through the nuclear β-actin [Actb] locus; Supp. Fig. 4I).

To further validate these results, we established a method for quantifying immunostained proteins and mtDNA in somatodendritic and axonal mitochondria of a fifth neuronal system: dentate gyrus neurons (DGN) in the hippocampus. This model is well-suited for spatially resolved analysis due to its well-defined regional anatomy and minimal intercompartmental distance, which ensures uniform tissue and staining quality (Fig. 3G-H). We co-stained the GFP-OMM tag with mtDNA and the mtDNA-associated protein Mitochondrial transcription factor A (Tfam), both of which are localized to mitochondrial nucleoids (Fig. 3I). Segmentation and quantification of mitochondrial regions further confirmed a significant reduction (0.25 ± 0.04, p = 0.001) in axonal mtDNA in DGN, accompanied by a significant but less pronounced decrease (0.64 ± 0.03, p = 0.0007) in Tfam. Notably, in samples with mtDNA and Tfam co-staining, the mtDNA-to-Tfam ratio – interpreted to predict mtDNA transcriptional activity (Brüser et al., 2021) – was also reduced (0.4 ± 0.09, p = 0.03) in axonal mitochondria, indirectly supporting active mtDNA expression as a somatodendritic mitochondrial feature (Fig. 3J).

### Axonal mitochondria are impoverished

Next, we combined this immunofluorescence-based quantification approach in DGN with our compartmentalized MitoTag analysis to systematically compare the expression level of mitochondrial proteins in the axon and the somatodendritic compartment. To select suitable targets, we constructed a ranked list of the differentially expressed mitochondrial proteins in our integrated MitoTag proteomes ordered from axonal to somatodendritic polarization (Supp. Fig. 5). We selected a panel of proteins distributed along the ranked list and directly compared their abundance in somatodendritic and axonal DGN mitochondria. As expected, proteins predicted to be somatodendritic showed clear enrichment in the corresponding compartment. Surprisingly, the proteins predicted to be axonally polarized were not more abundant in axonal mitochondria. Rather, they were mildly depleted in the axon or at best unchanged (Fig. 4A-C). The apparent contradiction between the proteome and microscopy data can be explained technically: our proteomic dataset is purified – first by immunocapture and then by *in silico* normalization – to contain exclusively mitochondrial proteins, yielding relative quantification within the mitochondrial proteome (i.e., it indicates the fraction of a mitochondrial protein over the total mitochondrial proteome). In contrast, immunofluorescence quantifies protein expression based on integrated intensity rather than fractional protein content, allowing a direct comparison of protein abundance in the two compartments. Nonetheless, the correlation between the log_2_ fold change from immunofluorescence and the proteomic rank was strong and statistically significant (Pearson r = 0.67, p = 0.01; Fig. 4D).

**Fig. 4:**
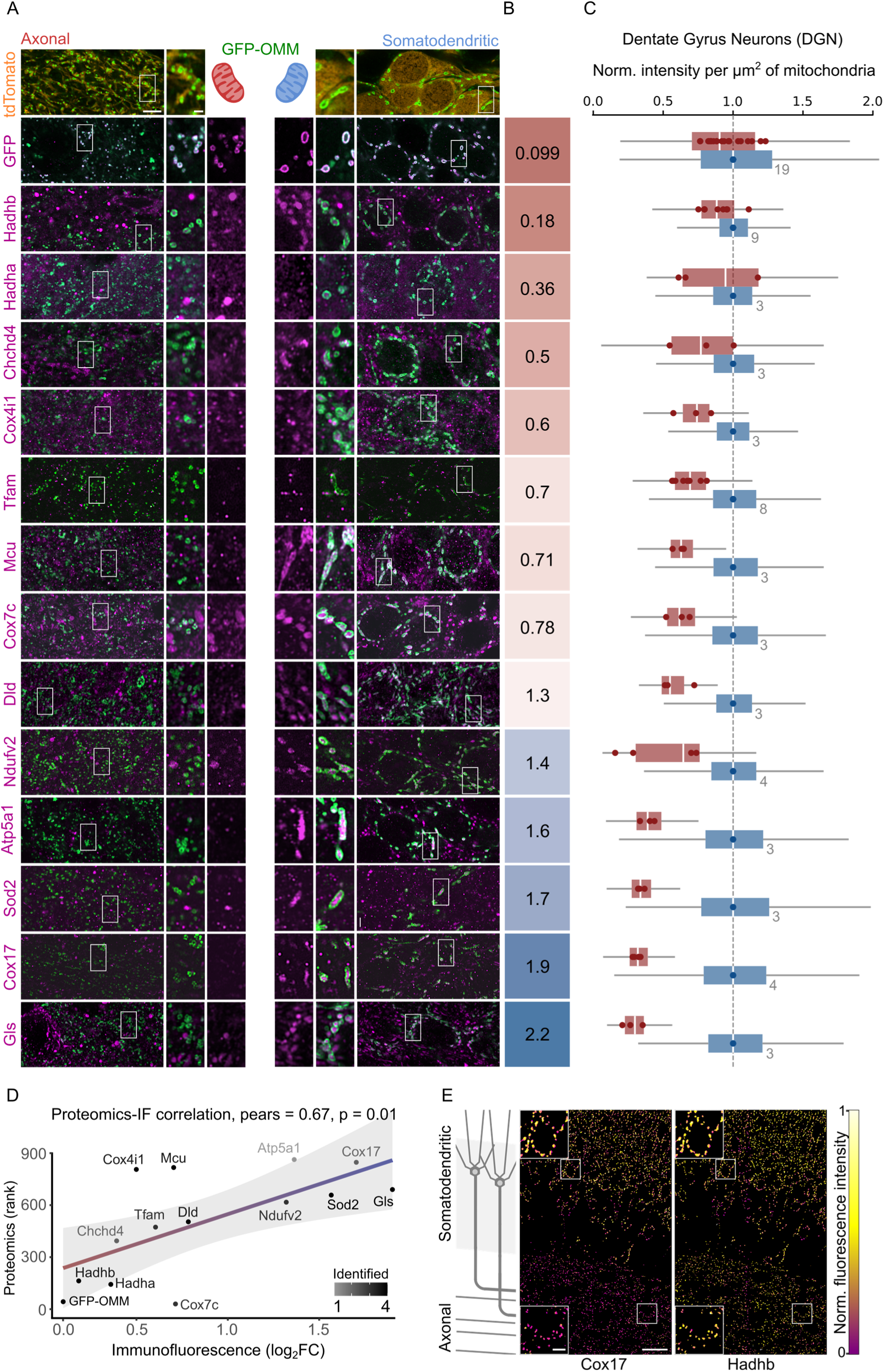
Immunofluorescence validation of rank-ordered mitochondrial proteins. **A.** Representative Airyscan immunofluorescence images of proteins with varying axon-to-soma ranks. Left panels: DGN axons showing co-localization between the protein of interest (POI) and axonal mitochondria. Right panels: co-localization with somatodendritic mitochondria. Anti-GFP: green, anti-POI: magenta, tdTomato: orange. Scale bar: upper left 5 µm (left) and 1 µm (right). **B.** Average axon-to-soma (Ax/So) ratios of POI intensity per µm^2^ mitochondrial cross-section area. **C.** Distribution of POI intensity per µm^2^ mitochondrial cross-section area. Points represent the median value per mouse (number of mice indicated in grey). Tfam box plot as in Fig. 3J. **D.** Correlation between immunofluorescence Ax/So ratios (panel B) and axon-to-soma protein ranks from all proteomics datasets. Pears: Pearson’s correlation coefficient. **E.** Heat map showing spatial distribution of Cox17 and Hadhb within segmented mitochondria. Scale bar 25 µm (right), 5 µm (left, detail).

Consistent with the functional data, mtDNA as well as oxidative phosphorylation proteins, including Cox17, Atp5a1, and Ndufv2, exhibit somatodendritic enrichment. Notably, Cox7c and Cox4i1 deviate from this trend; Cox7c possesses a high axonal rank (percentile ∼3), while Cox4i1 is highly somatodendritic (percentile ∼92), which does not correlate with the immunofluorescence ratio (Fig. 4D). By contrast, fatty acid oxidation proteins, such as Hadha and Hadhb, appear evenly distributed across compartments. So, while no single protein offers an isolated marker for axonal mitochondria, inversely polarized proteins (such as Cox17 and Hadhb) can provide direct visual confirmation of intraneuronal mitochondrial diversity, e.g., in DGN (Fig. 4E).

Taken together, these data suggest that axonal mitochondria are not enriched in specific pathways or proteins, but rather “impoverished”, i.e., characterized by reduced levels of many mitochondrial proteins, mtDNA, and several classical mitochondrial functions that somatodendritic mitochondria possess. At the same time, some pathways, such as fatty acid oxidation, are preserved. This renders axonal mitochondria, despite their broad impoverishment, relatively enriched and proficient in these functions, and comparable to somatodendritic mitochondria.

### Local translation maintains short-lived lipid-metabolism proteins in axonal mitochondria

Local translation represents a plausible mechanism underlying compartment-specific mitochondrial diversity (Hafner et al., 2019). It requires the presence of mRNA in the axon, which allows the local supply of proteins. To investigate this, we isolated ribosomes from our CCN neuronal population in a RiboTag mouse (Sanz et al., 2019) and sequenced the associated mRNAs, providing a snapshot of the transcripts actively engaged in translation within each compartment (Fig. 5A). The results were highly reproducible across four biological replicates (Fig. 5B).

**Fig. 5:**
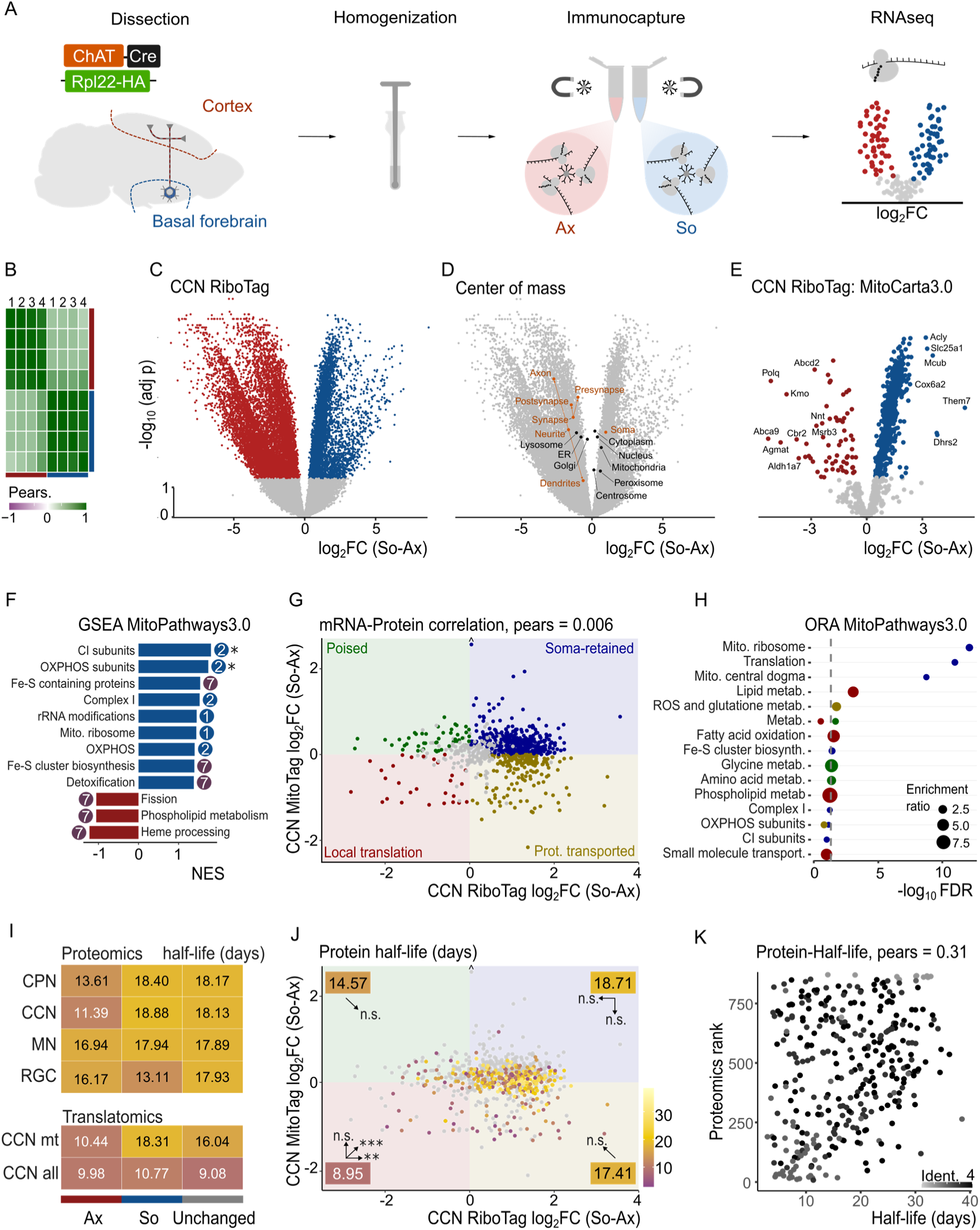
Local translation as a potential mechanism for compartment-specific diversity. **A.** Schematic of immunocapture of cell type- and compartment-specific ribosomes using RiboTag in CCN. From left: dissection of the brain regions of interest, mechanic homogenization, immunocapture of compartment-specific ribosomes with magnetic beads, RNA sequencing followed by data analysis **B.** Correlation heat map of biological replicates. **C.** Differential expression analysis (volcano plot) of all identified genes. Moderated t-test (limma); significance defined as FDR < 0.05. Genes are color-coded as somatodendritic (blue), axonal (red), or non-significant (grey). **D.** Center-of-mass projections of gene groups annotated for organelle and subcellular compartment localization. **E.** Differential expression analysis (volcano plot) of MitoCarta3.0 genes. Moderated t-test (limma); significance defined as FDR < 0.05. Genes are color-coded as in panel C. **F.** Gene set enrichment analysis using MitoPathways3.0. Somatodendritically (blue) and axonally enriched (red) pathways are labeled with their corresponding pathway groups from Fig. 2H. Asterisks indicate significance (FDR < 0.05). **G.** Correlation plot of translatomics (panel E) and proteomics (Fig. 2C). Categories are color-coded: blue, red, yellow, and green if significant in at least one test; grey indicates non-significant. **H.** Overrepresentation analysis of gene categories defined in panel G. Vertical line indicates significance threshold (FDR < 0.05). **I.** Heat map of protein half-life by compartment across all datasets. CCN mt: includes only MitoCarta3.0 genes; CCN all: includes all genes. **J.** Same data as in panel G, color-coded by protein half-life. Category averages are indicated in corners. One-way ANOVA with post hoc Tukey test: p < 0.05 *, < 0.01 **, < 0.001***; n.s., not significant. **K.** Correlation between proteomics rank (Fig. 4D) and protein half-life. Points are color-coded by the number of proteomes in which the protein was identified. Pears.: Pearson’s correlation coefficient.

Differential expression analysis in the two compartments revealed a substantial number of differentially localized transcripts: 8862 axonal and 4978 somatodendritic (Fig. 5C). We used UniProt annotations to group proteins by predicted cellular location and mapped the center of mass of these protein groups in the compartment-specific translatome volcano plot. This showed that proteins with a predicted somatic localization were preferentially translated on the somatodendritic side, while neurite- and synapse-associated gene products are expressed in the axon (Fig. 5D). While some dendritic and postsynaptic annotations also centered on the axonal side, albeit less strongly than axonal and presynaptic genes this likely reflects shared protein functions across these domains, such as the neuritic cytoskeleton or ion channels. Organelle annotations also supported compartmentalization: somatodendritic transcripts were enriched for gene products localized to the nucleus, centrosome, peroxisome, and mitochondria (consistent with the observed proteomic impoverishment of axonal mitochondria). In contrast, axonal transcripts contained lysosomal and endoplasmic reticulum annotations, while cytoplasm and Golgi apparatus annotations were equally distributed (Fig. 5D).

Differential expression and gene set enrichment analysis (GSEA) of mitochondrial transcripts revealed a partial overlap with our proteomic data. We found 65 axonal and 586 somatodendritic enriched mitochondrial transcripts (Fig. 5E). Somatodendritic mRNAs were enriched for oxidative phosphorylation and metabolism of metal and cofactor pathways, while axonal transcripts showed a trend towards mitochondrial fission, consistent with their smaller size (Lewis et al., 2018), phospholipid metabolism, and heme-processing (Fig. 5F). However, direct correlation analysis between the expression levels in proteomic and transcriptomic datasets yielded no significant relationship (Pearson r = 0.006, p = 0.87; Fig. 5G), in line with previous reports showing minimal correlation between mRNA and protein levels, particularly across independent samples (Edfors et al., 2016; Koussounadis et al., 2015). To obtain a less-fine grained sense of a possible contribution of local translation to compartmental mitochondrial diversity, we classified genes into five categories: (1) 397 soma-retained genes, for which protein and mRNA localized to the somatodendritic compartment, (2) 28 locally translated genes (both, mRNA and protein present in the axon), (3) 191 protein-transported genes (mRNA restricted to soma, protein present in axon), (4) 47 “poised” genes (mRNA present in axon, but protein underrepresented in this compartment), and (5) 229 unchanged genes (no significant compartmental difference). Overrepresentation analysis revealed that translation of genes related to mtDNA expression pathways was confined to the soma, while lipid metabolism and fatty acid oxidation-related genes were enriched among locally translated mRNAs in the axon. Reactive oxygen species (ROS) and glutathione metabolism were associated with protein-transported genes, and amino acid and glycine metabolism fell into the “poised” category (Fig. 5H). Interestingly, glycine metabolism proteins were specifically detected as enriched in the axons of CPN, but not in CCN, which could indicate a cell-type-specific role of axonal mitochondria in the metabolism of neurotransmitters (Casanova et al., 2023).

To further contextualize these findings and explore the converse role of mitochondrial protein degradation, we integrated our datasets with published protein half-life data (Fornasiero et al., 2018). On average, axonal proteins tended to have shorter half-lives than their somatodendritic counterparts (Fig. 5I). In particular, the gene group we classified as locally translated proteins exhibited significantly shorter half-lives than those in other categories (Fig. 5J), suggesting a need for continuous protein synthesis. This difference was statistically significant when compared to soma-retained (p < 0.001), unchanged (p = 0.02), and protein-transported (p = 0.002) categories, but not when compared to “poised” genes (p = 0.32). Despite these trends for groups of proteins, the correlation between lifetime and protein compartmentalization is significant but rather weak (Fig. 5K), suggesting a complex interplay of various biosynthetic and proteostatic mechanisms to create the unique mitochondrial populations in a neuron’s distinct cellular compartments.

## Discussion

Our study provides a comprehensive characterization of compartment-specific mitochondrial diversity by comparing somatodendritic and axonal mitochondrial proteomes across a range of neuron types. Somatodendritic mitochondria are comparatively enriched in proteins involved in mtDNA maintenance, transcription/ translation, oxidative phosphorylation, metal/ cofactor metabolism, and – to a lesser degree – proteostasis and calcium handling. In contrast, axonal mitochondria are comparatively depleted in these pathways yet retain the ability to metabolize lipids and amino acids, suggesting a distinct metabolic tuning. Together, our findings support a model in which somatodendritic mitochondria are optimized for ATP production and biogenesis. Axonal mitochondria, in contrast, are adapted to support distal metabolic homeostasis around metabolites that are critical, e.g., for neurotransmission, while abandoning a broader set of functional capabilities.

Beyond providing insights into the functional potential of mitochondria in the polarized cellular landscape of a neuron, our study also paints an unexpected picture of axonal mitochondria. Despite the initial impression from our proteomic “enrichment” scores, our data are less supportive of a model in which these mitochondria contain unique proteins or pathways that define them as “axonal”. Rather, our immunofluorescence-based comparison of mitochondrial protein levels in the hippocampus reveals that axonal mitochondria are differentially depleted in certain features of somatodendritic mitochondria, while retaining others. Therefore, we conclude that intraneuronal diversification of mitochondria is in part driven by the selective “impoverishment” of axonal mitochondria.

Such mitochondrial impoverishment can in principle be achieved through various mechanisms, likely involving complex and multifaceted processes. The two most plausible, non-exclusive possibilities are: (1) a sorting process based on mitochondrial dynamics in the soma, and (2) selective turnover in the axon, building on coordinated biogenesis and degradation.

The first possibility presumes a sorting mechanism that initially diversifies mitochondria and then directs molecularly specified mitochondria into the axon. This model predicts a molecular boundary at the junction between the soma and the axon. In this scenario, two distinct mitochondrial populations would exist near the axon hillock: One, which already carries a diminished proteome, would selectively enter the axon and constitute the anterogradely transported motile fraction. A molecularly more complete subpopulation would be selectively retained and remain resident in the soma or ascend into the dendrites. Past electron microscopy studies of the axon hillock have reported no apparent specializations in mitochondrial morphology (Palay et al., 1968); more recently, a mitochondrial clustering pattern in the axonal initial segment has been identified in a small percentage of neurons (Tjiang & Zempel, 2022). Moreover, there is recent precedent for the local functional and molecular specialization of mitochondria for bioenergetic versus biosynthetic roles even in compact cells, such as fibroblasts, utilizing a fission-based segregation (Ryu et al., 2024). Combined with compartment-specific fission processes that, e.g., have been described for dendritic mitochondria *in vivo*, a sorting-based specialization is conceivable (Virga et al., 2024). Furthermore, the selective sorting of mitochondria into dendrites versus axons has been previously ascribed to the differential engagement of TRAK/Milton transport adaptors, albeit without demonstrating other differences in the selected organelle pools (van Spronsen et al., 2013). So far, we have not yet observed a sharp distinction at the soma-to-axon boundary, but rather a progressive change in mitochondrial protein abundance along the axon (Supp. Fig. 3).

This favors the second model, based on progressive degradation, which implies a gradual rather than sharp molecular divergence of mitochondria along the axon. A simple “run-down” model, where axonal preservation is determined by long protein lifetimes, still seems unlikely. Here, protein lifetimes and functions would need to be coupled across hundreds of proteins to create a functionally meaningful outcome. Our bioinformatics correlation of published protein lifetimes versus axonal preservation does not reveal a monofactorial relationship (Fig. 5K). More likely – and supported by our combined proteome/translatome analysis (Fig. 5G-K) – is a complex relationship between local protein synthesis and degradation with a bias toward the local translation of short-lived proteins. This interplay of lifetime and local replenishment depletes unstable proteins in axonal mitochondria unless they are selectively preserved through local synthesis. Recent work has directly demonstrated this for the short-lived Pink1 protein, as axonally transported mitochondria carry the Pink1 mRNA tethered by specific RNA-binding proteins (Harbauer et al., 2022). There are, however, also examples of multifactorial stabilization: Cox7c, one of the most axonally preserved proteins located at percentile ∼3 in our proteomic dataset (Fig. 4A, D), has an extended half-life (Krishna et al., 2021), but also undergoes local translation in axons (Cohen et al., 2022).

Despite such examples, the extent to which local translation controls the compartment-specific mitochondrial proteome remains unclear, in part because correlated analyses of local proteomes and translatomes in fully differentiated neurons have rarely been reported. *In vitro* systems, such as microfluidic chambers, enable such studies by physically separating axons from somata; however, they do not always represent a mature and homeostatic state of neurons. Such *in vitro* investigations have reported an enrichment of mitochondrial transcripts, including oxidative phosphorylation components, in axons (De Pace et al., 2024; Maciel et al., 2018; Minis et al., 2014). Notably, a recent report, while concurring that ATP generation depends on local translation, also suggests that this process is still inefficient in axons when compared to somata (Zaninello et al., 2023). Other studies go a step further and suggest that axonal mitochondria do not use oxidative phosphorylation to produce ATP (Hirabayashi et al., 2024). When examined *in vivo*, the extent of the role of local translation in mitochondrial maintenance becomes less clear (Farias et al., 2020; Jung et al., 2023). This contrast suggests that while *in vitro* models provide important mechanistic insights, they may not fully capture the dynamic and context-dependent nature of local translation *in vivo*. Additionally, selective and local degradation, either through proteostasis, release and degradation of mitochondria-derived vesicles, or piecemeal mitophagy (Misgeld & Schwarz, 2017) likely further refine the resulting compartment-specific mitochondrial proteome, while also contributing to the characteristic compact shape of axonal mitochondria (Fischer et al., 2018).

Whatever the mechanism, a clear outcome of our study is that the differences between somatodendritic and axonal mitochondria are rather broad. We have not identified any unique protein that is selectively shipped into the axon and could, in isolation, be used to define axonal mitochondria. There could be some technical reasons for this, as our somatodendritic proteomes have a variable degree of axonal contamination depending on the length of the axon stretch and its local collaterals within the somatodendritic isolation. This would reduce the fold change of such single proteins and might make them difficult to detect as differentially expressed. Still, our results are in line with the fact that, to our knowledge, no truly “axon-specific” mitochondrial proteins have been described. Of the two TRAK isoforms, which have been reported to show some neuritic polarity (van Spronsen et al., 2013), we only detected TRAK1, which was relatively preserved in axons (percentile ∼33). However, due to the transient association of TRAKs with mitochondria, the capture and assignment of these proteins in our workflow should be interpreted with caution. Additionally, two other mitochondrial proteins, Syntaphilin (Snph) and Mitochondrial fission factor (Mff), perform well-characterized roles specifically in axonal physiology. Snph regulates mitochondrial distribution by anchoring mitochondria to microtubules, while Mff controls mitochondrial size not only at the axon entry point, but also maintains it along the axon shaft (Chen & Sheng, 2013; Lewis et al., 2018). However, despite these intraaxonal functions, these proteins do not appear to be strongly enriched in axons (e.g., Kang et al., 2008). Indeed, our proteomic data position Snph at a percentile rank of ∼29 and Mff at ∼95 in the axon-to-soma ranked polarity list, suggesting mild enrichment or even depletion in axons.

Beyond insights into the molecular polarization of mitochondria, our data also point toward functional diversification. We probed this using *ex vivo* assays on somatodendritic and axonal mitochondrial isolations. We chose this reductionist approach deliberately, rather than *in situ* methods, e.g., biosensors (Breckwoldt et al., 2014), as the local cellular environment or the functional state of the host neuron is very likely to affect the specific functional capabilities of a mitochondrion. Indeed, molecular diversity could act as much to maintain the key functionality of mitochondria locally stable as to adapt the organelle to specific local needs in their cellular context. For example, molecular diversity might compensate for organelle-intrinsic differences, such as shape heterogeneity (e.g., the altered ratio of different mitochondrial compartments in more spherical axonal versus more elongated dendritic mitochondria) or exogenous factors (e.g., organelle contacts or local signaling).

Our proteomic analysis predicted functional features that we could confirm using isolated mitochondria from the two compartments. For instance, the higher efficiency of somatodendritic mitochondria to utilize classical complex I and II substrates, compared to a comparatively preserved ability of axonal mitochondria to support fatty acid oxidation (Fig. 3A-B), chimes with the fact that oxidative phosphorylation as a pathway has its center of weight towards the soma, while lipid metabolism is a feature that axonal mitochondria preserve (Fig. 2H-L). Our integration of proteomics and translatomics data suggests local translation of lipid metabolism enzymes (Fig. 5H). This might allow axonal mitochondria to remain bioenergetically competent for specific substrates, compensating the partial depletion of the oxidative phosphorylation machinery by utilizing more energy-efficient fuels such as fatty acids.

These results are in line with established and emerging evidence: Early computational models proposed the postsynaptic compartment to be more energy-consuming than axon terminals (Attwell & Laughlin, 2001; Harris et al., 2012). Prior work using synaptosome proteomics indeed demonstrated low levels of proteins related to oxidative phosphorylation (Stauch et al., 2014). A current preprint similarly demonstrates, through imaging and single organelle characterization, that axonal mitochondria are scarce in mtDNA (Hirabayashi et al., 2024). Whether this implies a general inefficiency of axonal mitochondria in contributing ATP to axon and presynapse or rather hints at further local diversity and functional plasticity remains to be demonstrated (Rangaraju et al., 2014).

At the same time, several lines of evidence are converging on a hitherto unanticipated role of axonal mitochondria in lipid metabolism. Here, our data is in line with parallel work on synaptosome preparations (Antunes et al., 2025). However, synaptosomes could include astrocytic mitochondria, which are also enriched in fatty acid oxidation enzymes and capable of lipid, ketone, and amino acid metabolism (Fecher et al., 2019; Völgyi et al., 2015), complicating the interpretation. These results are, however, supported by the insight that the DDHD2 lipase (which itself is not mitochondrial) can fuel beta-oxidation in axons. This energy source is essential for maintaining distal metabolic homeostasis and sustaining local ATP levels (Antunes et al., 2025; Kumar et al., 2025; Saber et al., 2024; notable other roles in presynaptic lipid use and plasticity have also been proposed; Akefe et al., 2024). In our analysis, Hadhb and Hadha – which encode the alpha- and beta-subunits of the mitochondrial trifunctional protein (MTP), involved in beta-oxidation – are among the most prominent axonal hits, ranking at percentile ∼19 and ∼16, respectively, in our axon-to-soma polarity list from proteomics and exhibiting the best axonal preservation in our immunostainings (Fig. 4A-C). Notably, loss-of-function mutations in Hadhb and Hadha are associated with axonal pathology of Charcot-Marie-Tooth disease and sensorimotor axonopathy (Balletto et al., 2025; Hong et al., 2013; Nadjar et al., 2020), underscoring the importance of lipid metabolism in axonal homeostasis.

Calcium signalling and buffering represent other key functions of axonal mitochondria. Interestingly, protein-level analysis of the mitochondrial calcium uniporter (Mcu) suggested strong somatodendritic enrichment (percentile ∼93), which was confirmed, albeit less markedly, by immunostaining (Fig. 4A-C) and is consistent with prior synaptosome-based studies (Stauch et al., 2014).

However, in our study, Mcu enrichment alone was not a good predictor for function, since axonal mitochondria sustained calcium uptake more effectively during prolonged stimulation (Fig. 3C-D). Notably, the functional properties of the MCUC are not only shaped by Mcu abundance, but also by the stoichiometry with regulatory components such as Micu1-3 and Emre, as well as post-translational modifications (Rodríguez-Prados et al., 2023; Y. Wang et al., 2019; Yamada et al., 2023). The literature provides strong evidence for a role of Micu3 at synapses (Ashrafi et al., 2020). In our data, Micu3 is the most abundant MCUC protein in axons after Mcu (Supp. Fig. 4C), and yet it is strongly somatodendritically enriched (percentile ∼97). Recent compartmentalized *in vitro* studies also support somatodendritic expression of Mcu and Micu3 with balanced stoichiometry (Jang et al., 2025). Indeed, stoichiometry may matter more here than expression levels. Our analysis of the ratios of MCUC components revealed a stable complex stoichiometry across compartments, with the exception of Emre, which was overrepresented in axonal mitochondria. Emre stabilizes the core MCUC, but importantly, also shapes its function by bridging the main channel to the Micu regulators (Sancak et al., 2013; Y. Wang et al., 2019), potentially contributing to the enhanced buffering performance observed during the saturation phase. Notably, in absolute terms, Emre also does not exhibit axonal preservation (percentile ∼60) and shows high neuron-to-neuron variability (neuron-type coefficient of variation; ntCV = 0.48), suggesting that calcium dynamics might be regulated with substantial cell-type and subcellular specificity (Ashrafi et al., 2020; Fecher et al., 2019).

Given such divergence, our study – while providing a first systematic view of the intraneuronal diversity of mitochondria *in vivo* – also has limitations. Beyond the caveats that proteomes should not be misunderstood as invariable determinants of function, we probably underestimate the full extent of organelle diversification. On the one hand, we focused on reporting differences shared by several neuronal subtypes, integrating data of varying quality due to the size of the profiled mitochondrial populations. While this approach provided robustness, it overlooks cell-type-specific diversification, e.g., related to neurotransmitter type, which could still be detectable within our dataset (Fig. 2B-E). On the other hand, the emerging literature on local biosynthesis and transfer of mitochondria, their enzymes or metabolites, and the specific roles that such processes play in sustaining specific neuronal compartments (Asadollahi et al., 2024; Chamberlain et al., 2021; Rangaraju et al., 2019), suggests that mitochondrial diversity in neurons is even more nuanced than our study shows. Probably, local niches in neurons – at the pre- and postsynapse, near nodes of Ranvier and branch points – have their own functions and hence molecular specializations in mitochondria. These subcompartments are averaged together in our study. Therefore, we believe that further detailed investigations, not only of the composition, but also of the function of these various organelle pools *in situ*, will shed light on the even greater diversity that empowers neuronal mitochondria.

## Methods

### Animals

All animal experiments were approved by the responsible regulatory authority in Munich, Germany (Regierung von Oberbayern) or the Institutional Animal Care and Use Committee of Duke University. Mice (Mus musculus, C57BL/6 mixed N/J background; both male and female) were housed under a 12-hour light-dark cycle with ad libitum access to food and water. All experiments were conducted during the light phase. C57Bl/6J mice (Jackson Labs, 000664) were used as wild-type (WT) controls. To generate cell-type-specific MitoTag mice, the reporter strain (GFP-OMM^fl/fl^; Jackson Labs, 032675; Fecher et al., 2019) was crossed with various Cre-driver lines: Rbp4-Cre (031125-UCD; Mutant Mouse Resource and Research Center, MMRRC; Gong et al., 2003), ChAT-Cre (Jackson Labs, 006410; Rossi et al., 2011), or Vglut2-Cre (Jackson Labs, 028863; Vong et al., 2011). A tdTomato allele (Jackson Labs, 007914, Madisen et al., 2010) was also included in some Rbp4-Cre and ChAT-Cre mice for imaging purposes. Analogously, cholinergic-specific RiboTag mice were generated by crossing Rpl22-HA^fl/fl^ mice (Jackson Labs, 011029; Sanz et al., 2019) with the ChAT-Cre line. Ndufs4 knock-out mice (Jackson Labs, 027058; Kruse et al., 2008) were crossed with Vglut2-Cre; GFP-OMM^fl/fl^ for the methodological validation.

### Antibodies

The following primary antibodies were used, at the indicated maximum working dilutions, for immunofluorescence (IF) or Western blot (WB) analyses, either with heat-induced epitope retrieval (HIER) or without retrieval. HIER was performed in EDTA buffer (1 mM EDTA, pH 8.0; Sigma, E6758) supplemented with 0.05% Tween-20 (P1379; Sigma): rabbit polyclonal anti-ATP5A1 (1:100 IF; ProteinTech, 14676-1-AP), rabbit polyclonal anti-CHCHD4 (1:50 IF; ProteinTech, 21090-1-AP), rabbit polyclonal anti-COX4 (1:1000 WB; Abcam, ab16056), rabbit polyclonal anti-COX4I1 (1:50 IF; Abgent, AP22111a), rabbit polyclonal anti-COX17 (1:100 IF; ProteinTech, 11464-1-AP), rabbit polyclonal anti-COX7C (1:50 IF; ProteinTech, 11411-2-AP; HIER), rabbit polyclonal anti-DLD (1:200 IF; ProteinTech, 16431-1-AP), mouse monoclonal anti-dsDNA (1:66 IF; Progen, 690014S), goat polyclonal anti-GFP (1:1000 WB; Abcam, ab5450), chicken polyclonal anti-GFP (1:900 IF; Abcam, 13970), rabbit polyclonal anti-GLS (1:50 IF; ProteinTech, 20170-1-AP), rabbit polyclonal anti-HADHA (1:50 IF; Abcam, ab54477; HIER), mouse monoclonal anti-HADHB conjugated to A594 (1:50 IF; ProteinTech, CL594-67967), rabbit polyclonal anti-MCU (1:50 IF; Sigma, HPA016480; HIER), mouse monoclonal anti-NDUFS4 (1:1000 WB; Santa Cruz Biotechnology, sc-100567), rabbit polyclonal anti-NDUFV2 (1:100 IF; ProteinTech, 15301-1-AP), rabbit polyclonal anti-SOD2 (1:50 IF; ProteinTech, 24127-1-AP), rabbit polyclonal anti-TFAM (1:100 IF; ProteinTech, 22586-1-AP), mouse monoclonal anti-TUBB3 conjugated to A647 (1:250 IF; BioLegend, 657406), mouse monoclonal anti-TUJ1 (1:500 IF; Fisher Scientific, MAB11905), mouse polyclonal anti-VDAC (1:1000 WB; EMD Millipore, AB10527). The following secondary antibodies against the appropriate primaries were used in different combinations: goat anti-chicken IgY A405 (1:500 IF; ThermoFisher, A48260), goat anti-chicken IgY A488 (1:500 IF; ThermoFisher, A32931), goat anti-chicken IgY A647 (1:500 IF; ThermoFisher, A21449), goat anti-rabbit IgG A647 (1:500 IF; ThermoFisher, A21245), goat anti-rabbit IgG A405 (1:500 IF; ThermoFisher, A31556), goat anti-mouse IgG A594 (1:500 IF; ThermoFisher, A11005). Secondary antibodies against the appropriate species conjugated to IRDye 800CW or 680RD (1:20000 WB; Li-Cor, #926-32212 and #926-68073) were used for WB experiments.

### Mitochondria immunocapture

Mitochondria were isolated from freshly dissected mouse tissues -including spinal cord, sciatic nerve, ocular tissues, basal forebrain/striatum, and cortex-by adapting a previously published protocol (Fecher et al., 2019). Tissue was collected from male and female Promoter-Cre/GFP-OMM mice across a broad age range (3-22 months). Mice were euthanized by intraperitoneal administration of tribromoethanol (Avertin) for RGC mice, or by isoflurane overdose for CPN, CCN, and MN mice. This was followed by transcardial perfusion with ice-cold PBS for ≤ 3 min. Dissections were carried out immediately post-perfusion in ice-cold PBS according to the region of interest: CPN mice (cortex and striatum), CCN mice (cortex and basal forebrain), MN mice (spinal cord and sciatic nerve), and RGC mice (retina and optic nerve). Tissues were weighed and placed in isolation buffer (IB; 220 mM mannitol, 80 mM sucrose, 10 mM HEPES, 1 mM EDTA, pH 7.4), supplemented with protease inhibitor (cOmplete™, EDTA-free; Roche, 11873580001) and 1% essentially fatty acid-free BSA (Sigma, A7030) at 20 mg tissue per mL IB. For retina and optic nerve, 600 µL of IB was added per sample. All steps were performed on ice or at 4 °C. Tissues were homogenized in IB using a Dounce glass homogenizer (Sigma-Aldrich, D9063): three strokes with a type A pestle for brain tissues, and five to six strokes for spinal cord, nerves and ocular tissues. Retina and optic nerve underwent an additional six strokes with a type B pestle. Samples were subjected to nitrogen cavitation at 10 mg/mL using a pressure vessel (Parr Instrument Company, #4635) at 800 psi for 10 min. Pressure was released slowly. Lysates were centrifuged at 600 × g for 10 min at 4 °C to remove nuclei and large debris. The supernatant was passed through a 30 µm pre-separation filter (Miltenyi Biotec, #130-041-407) and diluted 1:6 in immunocapture buffer (ICB; 137 mM KCl, 2.5 mM MgCl₂, 3 mM KH_2_PO_4_, 10 mM HEPES, 1 mM EDTA, pH 7.4) supplemented with protease inhibitor and 1% BSA. Retina and optic nerve filtrates were centrifuged at 13,000 × g for 3 min to precipitate the mitochondria and resuspended in 2 mL ICB total volume. Superparamagnetic microbeads conjugated with anti-GFP (Miltenyi Biotec, #130-091-125) or anti-TOM22 (Miltenyi Biotec, #130-127-693) antibodies were added: 25 µL for brain, spinal cord and sciatic nerve and 60 µL for retina and optic nerve. Samples were incubated at 4 °C on a moving plate for 90 min, or on a rotating wheel for 60 min (retina and optic nerve). LS columns (Miltenyi Biotec, #130-042-401) were equilibrated with 3 mL ICB on a QuadroMACS separator (Miltenyi Biotec, #130-090-976). Samples were loaded in 3 mL increments (1 mL for retina and optic nerve). Columns were washed with 4 × 4 mL ICB (retina and optic nerve), removed from the separator and eluted in 3 mL ICB using a plunger. Eluates were centrifuged at 12,000-13,000 × g for 10 min. Pellets were washed in IB without BSA, and centrifuged at 8,000 × g for 3 min (except for optic nerve and retina). Mitochondria were stored at −80 °C for proteomic, Western blot, and mtDNA quantification, or used fresh for functional assays (e.g. calcium uptake, mitochondrial respiration). Cre-expression of all animals used for immunocapture in this study was monitored using an immersion-fixed spared brain region from each sample. Mice with unexpected Cre-expression pattern (e.g., Cre leak into astrocytes or germline recombination) were excluded.

### Total protein quantification

Protein concentration was determined using the Pierce BCA Protein Assay Kit (Thermo Fisher Scientific, #23225), following the manufacturer’s instructions. Briefly, mitochondrial pellet resuspended in 50 µl IB without BSA, then diluted 1:10 in RIPA buffer. BSA standards (ranging from 2 mg/mL to 25 µg/mL) and samples were pipetted in duplicates into a transparent, flat-bottom 96-well plate (ThermoFisher). BCA working reagent was added to each well at a 1:21 ratio (sample to reagent). The plate was incubated at 37 °C for 30 min and absorbance was measured at 562 nm using a plate reader (CLARIOstar, BMG Labtech). Sample concentrations were calculated by interpolation from the standard curve.

### Sample preparation and LC-MS/MS

Mitochondrial pellets were lysed in RIPA buffer and processed using a modified single-pot, solid-phase-enhanced sample preparation (SP3) protocol (Hughes et al., 2014). Briefly, 10-15 µg of protein per sample were diluted 1:2 with water, supplemented with 10 mM MgCl_2_ and 25 units benzonase (Sigma-Aldrich) for DNA digestion (30 min, 37°C). Proteins were reduced with 15 mM DTT (30 min, 37°C) and alkylated with 60 mM IAA (30 min, 20°C). Samples were bound to 40 µg of a 1:1 mix of hydrophilic/hydrophobic Sera-Mag SpeedBeads (Cytiva) in 80% ethanol (30 min, RT), washed four times with 80% ethanol, and digested for 16 h with 187 ng each of LysC and trypsin (Promega, V5072). Supernatants were filtered (0.22 µm Spin-X, Sigma-Aldrich), dried by vacuum centrifugation, and peptide concentrations estimated using the Qubit protein assay (Thermo Fisher). For brain mitochondria, 350 ng of peptide per sample were loaded onto a 15 cm in-house packed C18 column (75 μm ID, ReproSil-Pur 120 C18-AQ, 1.9 μm, Dr. Maisch GmbH) using a nanoElute nano-HPLC (Bruker). Peptides were separated using a binary gradient of water and acetonitrile (0.1% formic acid) at 300 nL/min (0 min: 2% B; 2 min: 5% B; 52 min: 24% B; 60 min: 35% B; 70 min: 60% B) at 50°C. The system was coupled to a TimsTOF Pro mass spectrometer (Bruker) via a CaptiveSpray ion source. Data were acquired using diaPASEF with a TIMS ramp time of 100 ms, one full MS scan, and 26 isolation windows (27 m/z width) covering 350-1001 m/z. Each diaPASEF scan included two isolation windows with a cycle time of 1.4 s. For spinal cord and sciatic nerve mitochondria, 1.3 µg of peptides were separated on a C18 column (300 mm × 75 µm, ReproSilPur 120 C18-AQ, 1.9 µm; Dr. Maisch) using an Easy nLC-1200 nano-HPLC (Thermo Fisher Scientific) with a binary gradient of water/acetonitrile (0.1% formic acid): (0 min: 2.4% B; 2 min: 4.8% B; 92 min: 24% B; 112 min: 35.2% B; 121 min: 60% B). The system was coupled via a Nanospray Flex ion source (Thermo Fisher) and column oven (Sonation) to a Q-Exactive HF mass spectrometer (Thermo Scientific). MS1 spectra were acquired at 120,000 resolution (AGC target: 3E+6). The top 15 peptide ions were fragmented by HCD (resolution: 15,000; isolation width: 1.6 m/z; AGC target: 1E+5; NCE: 26%) with dynamic exclusion of 120 s. Retina and optic nerve mitochondria digests were dissolved in 12 µL of 1/2/97% trifluoroacetic acid/acetonitrile/water; 4 µL were injected into a 5 µm, 300 µm × 5 mm PepMap Neo C18 trap column (Thermo Fisher) in 1% acetonitrile for 3 min at 5 µL/min. The analytical separation was performed on a 2 µm, 75 µm × 250 mm EasySpray PepMap C18 column (Thermo Fisher) over 90 min at 0.3 µL/min and 35°C using Vanquish Neo UPLC. A 2-30% mobile phase B gradient was used (A: 0.1% formic acid in water; B: 0.1% formic acid in acetonitrile). Peptides were analyzed on a Q Exactive HF Orbitrap (Thermo Fisher) using positive electrospray ionization (2000 V, 275°C). MS1 scans were acquired at 120,000 resolution (m/z 375-1450; AGC: 3.0e6; RF lens: 30%; max injection time: 50 ms). MS/MS was performed in DDA mode with charge state filtering (2-5), monoisotopic peak detection, dynamic exclusion (20 s, 10 ppm), and HCD (collision energy: 30±5%; AGC: 1.0e5; max injection time: 100 ms).

### Protein identification and quantification

For brain mitochondria, label-free protein quantification was performed using DIA-NN v1.8 (Demichev et al., 2020) in library-free mode. The search was conducted against a Mus musculus UniProt database (one protein per gene; 21,845 entries; 2023), supplemented with a contaminant database (124 entries) and GFP-OMM. Trypsin/P was set as the protease, allowing up to two missed cleavages. Variable modifications included methionine oxidation and protein N-terminal acetylation; carbamidomethylation of cysteines was set as a fixed modification. Precursor and fragment ion m/z ranges were 350-1001 and 200-1700, respectively, with precursor charge states of 2-4. MS1 and MS2 mass tolerances were set to 15 ppm. Ion mobility tolerance was automatically adjusted. A 1% FDR threshold was applied for both peptide precursor and protein identifications. Match-between-runs and cross-run normalization were enabled. For spinal cord and sciatic nerve, data were analysed using MaxQuant v1.6.3.3 (Cox & Mann, 2008). A one protein per gene Mus musculus UniProt database (22,294 entries; 2019) was used. Trypsin was defined as the protease, allowing one missed cleavage. Variable modifications included methionine oxidation and protein N-terminal acetylation; carbamidomethylation of cysteines was set as a fixed modification. FDR for peptides and proteins was controlled to <1% using a target-decoy approach. The match-between-runs feature was enabled with a 1.5 min time window. Label-free quantification (LFQ) required at least two ratio counts of unique peptides. For retina and optic nerve mitochondria, Progenesis QI for Proteomics v4.2 (Waters Inc.) was used for alignment and peak area calculation. Peptide identification was performed using Mascot v2.5.1 (Matrix Science) against the reviewed UniProt mouse database (17,008 entries; 2019) supplemented with GFP-OMM. Mascot parameters included: precursor ion tolerance of 10 ppm, fragment ion tolerance of 0.025 Da, one missed cleavage by trypsin, fixed modification of cysteine carbamidomethylation, and variable modifications of methionine oxidation and deamidation of asparagine/glutamine. Proteins identified with ≥2 peptides (peptide FDR <0.5%, protein FDR <1%) were included. Peptide peak areas were normalized using Progenesis total ion intensity correction. Protein abundance was calculated as the sum of non-conflicting peptide intensities, averaged across two technical replicates.

### Bioinformatics analysis of proteomics data

Proteomics bioinformatics data analysis was performed in R statistical software 4.5.1 (R Core Team, 2013), and the full code and package versions are available. Mitochondrial proteins were filtered based on the Mouse MitoCarta3.0 (MitoScore2.0 > 1) database (Rath et al., 2021), with the addition of Snph and GFP-OMM. Label-free quantification (LFQ) intensities were normalized by dividing the LFQ value of each protein by the total mito-LFQ value in its respective sample. Log_2_ transformation was applied to all normalized LFQ values. For the CPN and CCN brain datasets, IC-GFP intensities were background-corrected by subtracting IC-TOM values (within the same tissue), as done previously (Fecher et al., 2019). This correction was not applied to MN and RGC datasets due to substantial differences in background composition. Log_2_ fold changes between soma and axon compartments were computed for each protein, and statistical significance was assessed using the *limma* linear modeling framework (Ritchie et al., 2015) and GSEA with WebGestaltR (Liao et al., 2019). For Supp. Fig. 2J, the same analysis pipeline was applied but considering exclusively non-mitochondrial proteins. The analysis is summarized in Supp. Fig. 5.

### Mitochondria respiration

Mitochondrial respiration was assessed with Oroboros 2k-Oxygraph following the Bioblast guide. Briefly, Oxygraph-2k chambers were washed 3 times 5 min and equilibrated with 2.2 mL of MiR05 buffer (110 mM sucrose, 60 mM lactobionate, 20 mM taurine, 20 mM HEPES, 10 mM KH_2_PO_4_, 3 mM MgCl_2_, 0.5 mM EGTA, 1 mg/ml BSA, pH 7.1) for at least 1h in saturated oxygen conditions. Once the readout was stable, 30-50 µg of isolated mitochondria were added to each closed chamber, followed by the injection of the different substrates and inhibitors (modified SUIT-017_O2_mt_D046: 2 mM malate and 0.5 mM octaenoylcarnitine, 2.5 mM ADP, 10 µM cytochrome c, 5 mM pyruvate, 10 mM succinate, 0.5 µM CCCP [repeated until maximum capacity was achieved], 0.5 µM rotenone, and 2.5 µM antimycin A). The specific flux was calculated using the Excel O2 analysis template from the manufacturer and subtracting the background oxygen consumption.

### Calcium uptake

Calcium uptake was assessed as described previously (Wettmarshausen & Perocchi, 2017) with some modifications: 30-50 µg of isolated mitochondria were resuspended in 90 µL of respiration buffer (137 mM KCl, 2.5 mM MgCl_2_, 3 mM KH_2_PO_4_, 10 mM HEPES, 5 mM succinate, 5 mM malate, 5 mM glutamate or 2.5 mM Pyruvate, 0.2% essentially fatty acid-free BSA, pH 7.4, supplemented with 100 nM CalciumGreen-5N; C3737; Thermo Fisher Scientific). Then 10 µL of either respiration buffer or 10 µM Ru360 (final well concentration; Sigma, #557440) was added for a final volume of 100 µL. Fluorescence (excitation 485/emission 530) was measured with a plate reader (CLARIOstar, BMG Labtech) in 7-10 s intervals for at least 12 min with the injection of 20 µM CaCl_2_ (final well concentration) in respiration buffer every 3 min at 25 °C. Uptake response was calculated as the inverse of the area under the curve (AUC^-1^) using trapezoidal integration.

### Digital PCR

mtDNA and nDNA concentrations were assessed using the QIAcuity One-Plate Digital PCR System (QIAGEN, 911000). First, DNA was extracted by diluting (1:10) 2 µL of isolated mitochondria in QuickExtract DNA Extraction Solution (Biozym, 101098) and incubating at 65°C for 6 min, followed by 98°C for 2 min and then left to cool down on ice. Digital PCR analysis was then performed using Taqman assays and the QIAcuity Probe PCR kit (QIAGEN, 250102); a 15 µl reaction mix was prepared containing: 1.5 µl of TaqMan primer assay mix, 5.0 µl of QIAcuity MasterMix, 3.5 µl of sterile H_2_O and 5.0 µl of lysed DNA sample. A total reaction-mix of 12 µl was pipetted into each well of the QIAcuity Nanoplate (8.5k 96-well, 250021). To stay within the range of the digital PCR system, the lysed mitochondria sample had to be further diluted in distilled water to the equivalent of 0.1 pg of protein for mtDNA quantification with mt-Nd1 (ThermoFisher Mm04225274_s1; 4331182) or Cytb targeting primers (ThermoFisher Mm04225271_g1, 4331182). nDNA was diluted to the equivalent of 10 pg of protein and detected using primers targeting the Actb nuclear locus (ThermoFisher Mm02619580_g1, 4331182). Parameters used for dPCR were QIAGEN Priming Profile Probe (RT-) PCR: cycling for 1 x 95⁰C for 2 min, followed by 3-step cycling 40 x at 95 ⁰C, 60 ⁰C and 72 °C for 15 s each, and a final step at 40 °C for 5 min. The green fluorescence channel (FAM) was imaged with an exposure time of 300 ms and a gain of 6. Data analysis was performed using the concentration of DNA copy number given by the QIAcuity Software Suite v3.1.0.0 (QIAGEN).

### Western blotting

For WB analysis, mitochondrial pellets (3-5 mg) were resuspended in RIPA buffer containing 50 mM Tris-HCl, pH 8.0, 150 mM NaCl, 0.1 mM EDTA and 1% Triton X-100, 0.25% Nonidet P-40, 0.1% SDS, as well as 1x Protease Inhibitor cocktail on ice for 30 min with frequent vortexing. Samples were mixed with 1x LDS sample buffer (Thermo Fisher Scientific, NP0007) and heated at 70°C for 10 min. The sample was loaded onto a NuPAGE 4-12% Bis-Tris gel (ThermoFisher, NP0323) and SDS-PAGE was performed at 50 V for 20 min and 100 V for 40 min in 1x NuPAGE MES SDS Running Buffer (ThermoFisher, NP0002). Following electrophoresis, proteins were transferred using a semi-dry method (XCell II Blot Module, Novex EI9051) to a PVDF membrane (Bio-Rad, #1620177) pre-activated with methanol (30 minutes) at 35 V for 45 min. The membrane was blocked with a 1:1 mixture of PBS-Tween (0.0125%) and Li-Cor Intercept Blocking Buffer (Li-Cor, #927-700001), then incubated overnight at 4°C with primary antibody. After 3 sequential 15-minute washes in PBS-Tween (0.0125%) on a shaker at RT, the blot was incubated with secondary antibody for 1 h at RT, the washing sequence repeated, and imaged using the Li-Cor Odyssey CLx system.

### Electron microscopy

For ultrastructural analysis, purified mitochondrial pellets from RGC MitoTag retinas and optic nerves obtained via mitochondria immunoprecipitation were fixed in a solution of 2% glutaraldehyde and 2% paraformaldehyde in 1x PBS. The specimens were then post-fixed in a solution of 1% osmium tetroxide in 0.1% cacodylate buffer, followed by dehydration with graded ethanol from 30%-100% and propylene oxide. A 1:1 propylene oxide:Embed 812 Resin mixture was infiltrated overnight under a vacuum. Pure Embed 812 Resin was then exchanged on the second day and samples were incubated at 65°C overnight. Ultra-thin sections were cut at 54-75 nm thickness (Leica EM CU7) and contrast-stained with 1% uranyl acetate, 3.5% lead citrate solution. Image acquisition was performed on a JEM-1400 transmission electron microscope (JEOL) using a Gatan ORIUS (1000) camera.

### Tissue preparation and immunohistochemistry

Mice were euthanized by intraperitoneal injection of tribromoethanol (Avertin) for retina and optic nerve collection, or by isoflurane overdose for brain, spinal cord, and sciatic nerve tissues. Retina and optic nerve were dissected immediately following euthanasia and fixed in 4% paraformaldehyde (PFA) in PBS for 90 min on ice. For the remaining tissues, animals were transcardially perfused with 4% PFA in PBS (for ∼30 min), after which brain, spinal cord, and sciatic nerve were dissected and post-fixed overnight at 4 °C in 4% PFA in PBS, then transferred to PBS for storage at 4 °C. Brains and spinal cords were embedded in 3% agarose in PBS and sectioned at 50 µm using a vibratome (Leica, VT1200S). Free-floating sections or whole sciatic nerve were stored in PBS at 4 °C, permeabilized with 5% CHAPS (Roth, #1479.3) in distilled water for 1 h at 37 °C, and, if required, subjected to antigen retrieval (30 min at 90 °C in the appropriate buffer). Sections were washed once in PBS and blocked for 1 h at RT in a buffer containing 10% normal goat serum (Millipore, NMM1582560), 1% BSA, and 0.5% Triton X-100 (Sigma-Aldrich, T8787) in PBS. For retina and optic nerve cryosections, tissues were cryoprotected sequentially in 15% and 30% sucrose in PBS for 24 h each, embedded in Tissue Freezing Medium (Triangle Biomedical), and cryo-sectioned at 16 µm on a Leica CM1950 cryostat. Sections were mounted on glass slides, rehydrated in PBS, and blocked in 3% normal donkey serum in PBS containing 0.3% Triton X-100. All sections were incubated with primary antibodies diluted in blocking buffer overnight at 4 °C either in well plates (vibratome sections) or humidity chambers (cryosections). For quantitative comparison of axon versus soma compartment staining, primary antibodies were added in excess at the indicated dilutions. Following three 10-min washes in PBS, sections were incubated with secondary antibodies for 2 h at RT, then washed again and mounted on glass slides (Epredia, AG00000112E01MNZ10) with coverslips (Roth, #1871) using Fluoromount-G™ mounting medium (Invitrogen, 00-4958-02). Secondary antibody-only controls were performed to assess nonspecific binding.

### Airyscan and confocal microscopy

Tissue slides were scanned on an upright Zeiss Airyscan 2 LSM 980 microscope with 10x/0.45 N.A. air and 63x/1.4 N.A. oil DIC objective (Immersol™ 518F; Zeiss, 33802-9010-000). Airyscan images were pre-processed with a linear Wiener filter (Airyscan processing) using the Zen v3.9 software (Zeiss). The same parameters were selected for quantitatively comparable images. Retina and optic nerve cryosections were imaged on a Nikon Eclipse Ti2 inverted confocal microscope controlled by Nikon NIS-Elements AR v5.21.03 software (Nikon). A 40x/1.3 N.A. oil objective and Nikon A1 confocal scanner system were used for image acquisition. For figure representation, images were processed in the open source software ImageJ/FIJI v2.14.0 (Schindelin et al., 2012) by linearly adjusting contrast and brightness to improve visibility (except when indicated otherwise in the figure legends to enhance intermediate contrast in non-quantitative visualizations). Adjustments were always kept equal and linear between images that are meant to be compared quantitatively (e.g., axonal and somatodendritic mitochondria stained with the same antibody; Fig. 3I, 4A)

### Image analysis

The quantitative image analysis was performed in Python v3.12.4 and the full code and package versions are available. Briefly, a total of 580 images of dentate gyrus neuron compartments were segmented using the anti-GFP fluorescence channel (mitochondria) with a self-trained, random forest-based pixel classifier (cross-validation scores > 0.98). To quantify the protein of interest, mean fluorescence intensity per µm^2^ of mitochondria was extracted from the segmented mitochondrial regions (amounting to a total area of 175986 µm^2^), preserving spatial distribution. Systematic background subtraction was followed by the calculation of the median intensity values per mouse and compartment. The median axon values were normalized to the median intensity of the soma images for each mouse to allow comparison across compartments.

### Ribosome immunocapture, RNA sequencing, and analysis

Brain regions were dissected, snap-frozen in liquid nitrogen and kept at -80 °C until the pull down of labeled ribosomes as described previously (Gavoci et al., 2025; Sanz et al., 2019). Briefly, tissues were homogenized (buffer in RNase-free water in mM: 50 TrisCl, pH7.5, 100 KCl (Sigma, #P9541), 12 MgCl_2_ (Sigma, #63068), 1% NP-40 (Roche, #11332473001), 1 X dithiothreitol (DTT; Sigma, #646563), 1 X protease inhibitors (Sigma, #11697498001), 1 mg/ml heparin (Sigma, #H3393), 100 µg/ml cycloheximide (Sigma, #C7698), RNAseOUT (ThermoFisher, #10777019). As previously described, the mRNA-ribosome complex was precipitated using a polyclonal HA-antibody (Sigma, #H6908) and Dynabeads Protein G (Life Technologies, #10004D). Ribosome-bound mRNA was isolated with RNeasy Plus Micro Kit (Qiagen, #74034) per the manufactureŕs instructions. Prior to sequencing, total RNA quantity and integrity were assessed using the Agilent 2100 Bioanalyzer with the RNA 6000 Nano Kit, ensuring RNA Integrity Numbers (RIN) greater than 8.5. Ribosomal RNA was subsequently depleted using Ribo-Zero rRNA Removal Kit. CCN mRNA was sequenced in pair-end using an Illumina HiSeq4000 Kit at a depth of ∼ 40 million reads per sample (Institute of Neurogenomics, Juliane Winkelmann, Helmholtz Munich, Germany). Analysis of the data was carried out in Bash and R. Briefly, the raw sequencing data (fastq files) were assessed for quality with FastQC v0.12.1 (Andrews, 2010), aligned to the mouse genome with STAR v2.7.10b (Dobin et al., 2013), and read counts extracted with featureCount v2.0.1 (Liao et al., 2014). The count matrix was then assessed for the correlation between replicates and differential localization of transcripts performed with the limma linear modeling framework (Ritchie et al., 2015) and GSEA using WebGestaltR (Liao et al., 2019).

### Statistics

Differential expression analysis was tested with moderated t-test using the limma modeling framework. GSEA and ORA were tested for significance with weighted Kolmogorov-Smirnov test and the hypergeometric test, respectively with GebWestalltR. Functional and quantifications of mtDNA and Tfam were subjected to one-sample t-test, the reported p-values in the text are corrected for multiple comparisons (FDR) within each experiment. All correlations were tested for significance with the Pearson’s product-moment correlation test. Statistical tests and results of the analysis of these data are also reported in figure legends and article text.

### Use of large language models

Large language models were used for support in literature search (Perplexity), and language editing to improve readability (ChatGPT5, M365 Copilot, Grammarly).

## Acknowledgements

We thank N. Budak and D. Steinmetz for animal husbandry, K. Kellermann for veterinary support, and Y. Hufnagel, M. Schetterer, and K. Wullimann for technical and administrative assistance. We are grateful to L. Godinho (TUM) for manuscript input and D. Hoerl (LMU) for guidance on image analysis. We thank the technical support and instrument access for bioenergetic measurements provided by J. Sailer, H. Zischka (Institute of Toxicology and Environmental Hygiene, TUM), and for dPCR by S. R. Kalluri, R. Silveri, and B. Hemmer (Department of Neurology, TUM University Hospital). We thank Juliane Winkelmann and the Helmholtz Center Munich facility for RNA sequencing. Mass spectrometry data acquisition was supported by the SyNergy Proteomics Hub and the Max Planck Institute of Biochemistry; high-resolution imaging by the SyNergy Microscale Hub.

A.M.P. was supported by the Alzheimer Doctoral Scholarship from the Hans-and-Ilse-Breuer Foundation. T.M.’s lab was funded by the Deutsche Forschungsgemeinschaft (Mi 694/8-1, Mi 694/7-1, Mi 694/9-1 A03-ID 428663564) and the ERC under the European Union’s Seventh Framework Program (FP/2007-2013; ERC Grant Agreement n. 616791). Additional support came from TRR 274/1 (projects B03, C02, Z01, Z02, ID 408885537) and a DFG instrumentation grant (INST95/1755-1 FUGG, ID 518284373). T.M., A.B.H., M.S.B. and S.F.L. received support from the Munich Center for Systems Neurology (SyNergy EXC 2145; Project ID 390857198) and M.S.B. also by the DFG (project ID: 450131873) and the DGM foundation (ID Le3/1). A.B.H. is further supported by the Max Planck Society, the Deutsche Forschungsgemeinschaft (TRR353 - ID 471011418, SPP2453 - HA 7728/4-1 ID 541742535), the European Union (ERC StG Project 101077138 — MitoPIP), and the Schram Stiftung (T0287/46550/2025). S.F.L. was also supported through the Bundesministerium für Forschung, Technologie und Raumfahrt (BMFTR, FKZ: 01ED2504) under the aegis of JPND. R.C. and S.M.G.’s labs at Duke were supported by a BrightFocus Foundation National Glaucoma Research award, as well as a National Eye Institute Core Grant EY005722 to Duke University and an unrestricted grant from Research to Prevent Blindness to the Duke Eye Center. A.d.R.M was supported by the Walter Benjamin Programme from Deutsches Forschungsgemeinschaft (ID: 553397020), P.G-D. by the Alexander von Humboldt Research Fellowship, L.T. by the EMBO Fellowship (no. EMBO ALTF 108-2013), L.O. by the Monika and Walter Neupert Foundation and was a member of the International Max Planck Research School for Biological Intelligence, and M.A.A. by the DAAD-TEV-Master’s degree scholarship and was a student of the MSc program ‘Biomedical Neuroscience’ supported by the Elite Network Bavaria (ENB). A.M.P., L.O., P.P., A.G., and C.F. were also members of the Graduate School of Systems Neuroscience (GSN-LMU).

## Author contributions

A.M.P., L.S.C.L., R.C., S.M.G., and T.M. conceptualized the project and experiments. T.M., S.M.G., and R.C. supervised, administered, and provided infrastructure and funding for the project. A.M.P. and L.S.C.L. performed mouse breeding (with M.S.B.’s and H.M.B.’s support) and mitochondrial isolations (with contributions from P.G-D., P.P., and H.M.B.). A.M.P. performed Airyscan microscopy (with contributions from M.A.A.). L.S.C.L. performed confocal and electron microscopy (with contributions from Y.H.). L.O. and A.M.P. completed the dPCR experiments. A.d.R.M. and A.M.P. performed the calcium uptake assays. C.F. and L.T. established and characterized MitoTag mouse lines and isolation protocols. A.G. and P.G-D. performed the RiboTag experiments, while S.M., H.M.M., A.B.H., S.F.L., N.P.S., and V.Y.A. performed and contributed to the LC-MS proteomics. A.M.P. performed the bioinformatics and image analysis.

---

**Supp. Fig. 1:**
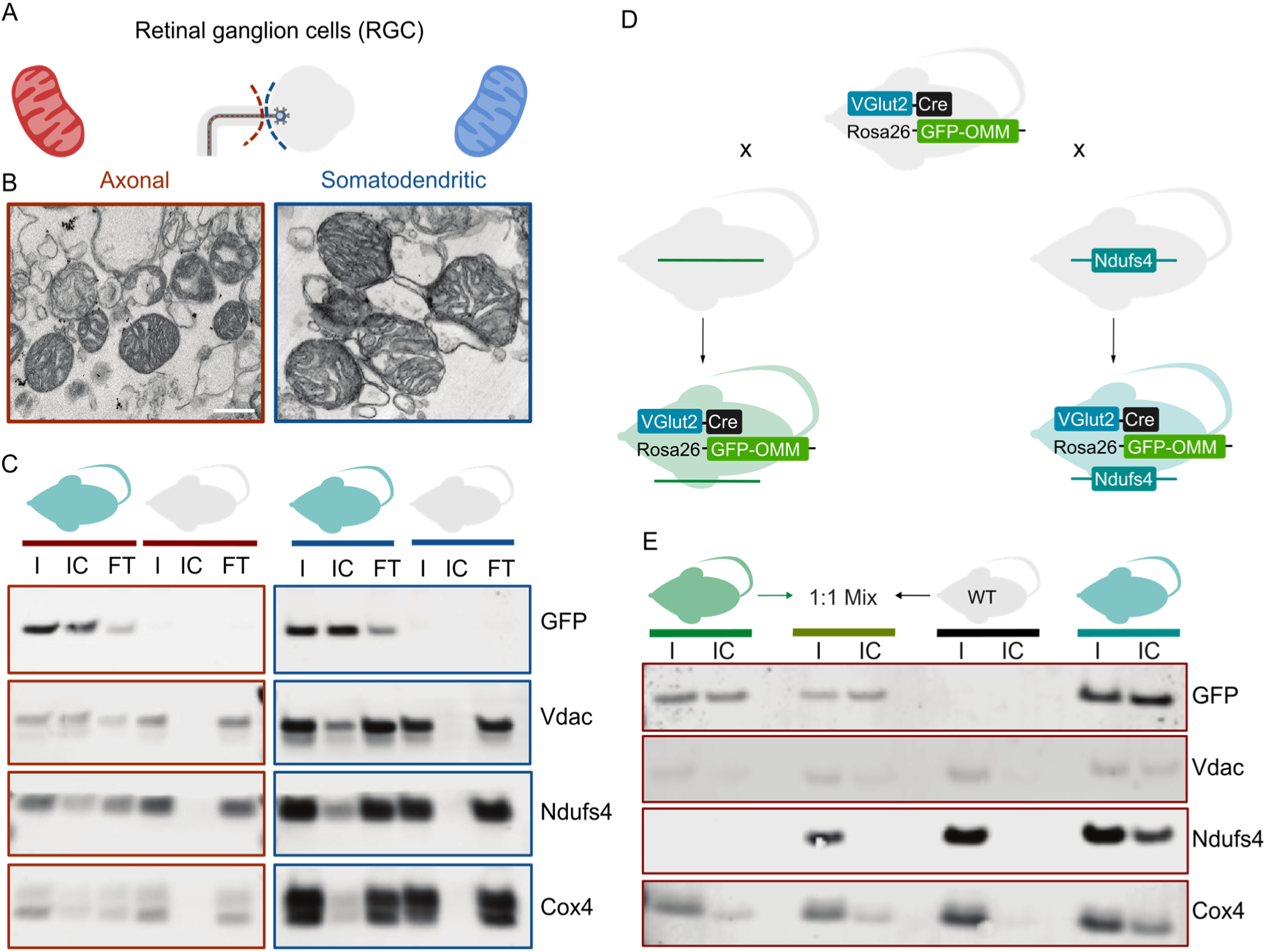
Isolation of compartment-specific RGC mitochondria using MitoTag. **A.** Schematic of RGC anatomy showing the somatodendritic compartment in the retina and the axonal compartment in the optic nerve. **B.** Electron micrograph of immunocaptured mitochondria from RGC compartments. Scale 500 nm. **C.** Western blot of immunocaptured mitochondria probed for GFP-OMM and endogenous mitochondrial proteins (Vdac, Ndufs4, and Cox4) from RGC-MitoTag mice (VGlut2-Cre/GFP-OMM, turquoise) and wild-type mice lacking the MitoTag (grey), from axonal (red) and somatodendritic (blue) isolates. **D.** Schematic of the generation of the VGlut2-Cre/GFP-OMM mouse line with Ndufs4 knockout (green mouse). **E.** Western blot of immunocaptured axonal mitochondria probed for GFP-OMM and mitochondrial proteins (Vdac, Ndufs4, and Cox4) from wild-type mice (grey), RGC (VGlut2-Cre: GFP-OMM)-Ndufs4-knockout (green), a 1:1 mix of these two (olive green), and RGC (VGlut2-Cre: GFP-OMM)-Ndufs4-wild-type (turquoise). I: input; IC: immunocaptured; FT: flow-through.

**Supp. Fig. 2:**
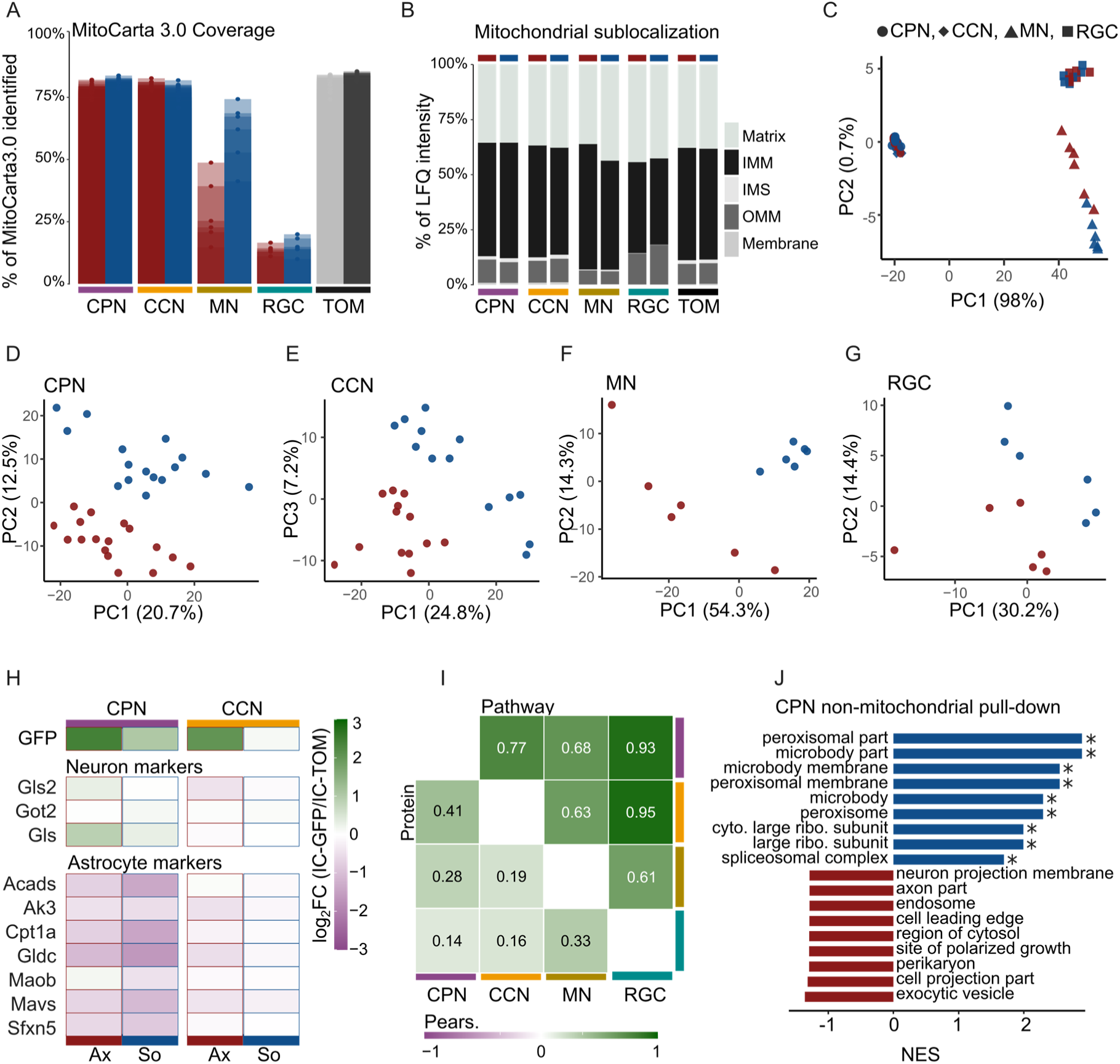
Evaluation and quality control of somatodendritic and axonal proteomes. **A.** Coverage of the MitoCarta3.0 database for neuron lines and IC-TOM. Bars from different biological replicates are superimposed. Axonal: red; somatodendritic: blue; striatal: grey; cortical: black. Neuron-types and IC-TOM (immunocapture-Tom22) are represented by distinct colors. **B.** Sub-organelle localization of mitochondrial proteins shown as percentages of total mitochondrial label-free quantification (LFQ) intensity. IMM: inner mitochondrial membrane; IMS: intermembrane space; OMM: outer mitochondrial membrane; Membrane: unassigned mitochondrial membrane. **C-G.** Principal component analysis of compartment-specific mitochondrial proteomes: C, all samples combined; D, CPN; E, CCN; F, MN; G, RGC. **H**. Heat map of protein ratios from cell type-specific immunocaptured mitochondrial proteomes (IC-GFP: immunocapture-GFP) over background tissue (IC-TOM). Analyzed proteins are validated neuronal or astrocytic markers (Fecher et al., 2019). **I.** Pearson correlation matrix of somatodendritic-to-axonal (So/Ax) ratios across neuronal types. Lower left triangle: protein log_2_ fold-changes; upper right triangle: pathway normalized enrichment scores. **J.** Gene set enrichment analysis of non-mitochondrial proteins from CPN using Gene Ontology Cellular Component. Somatodendritic pathways are shown in blue; axonal pathways in red. * indicates FDR < 0.05.

**Supp. Fig. 3:**
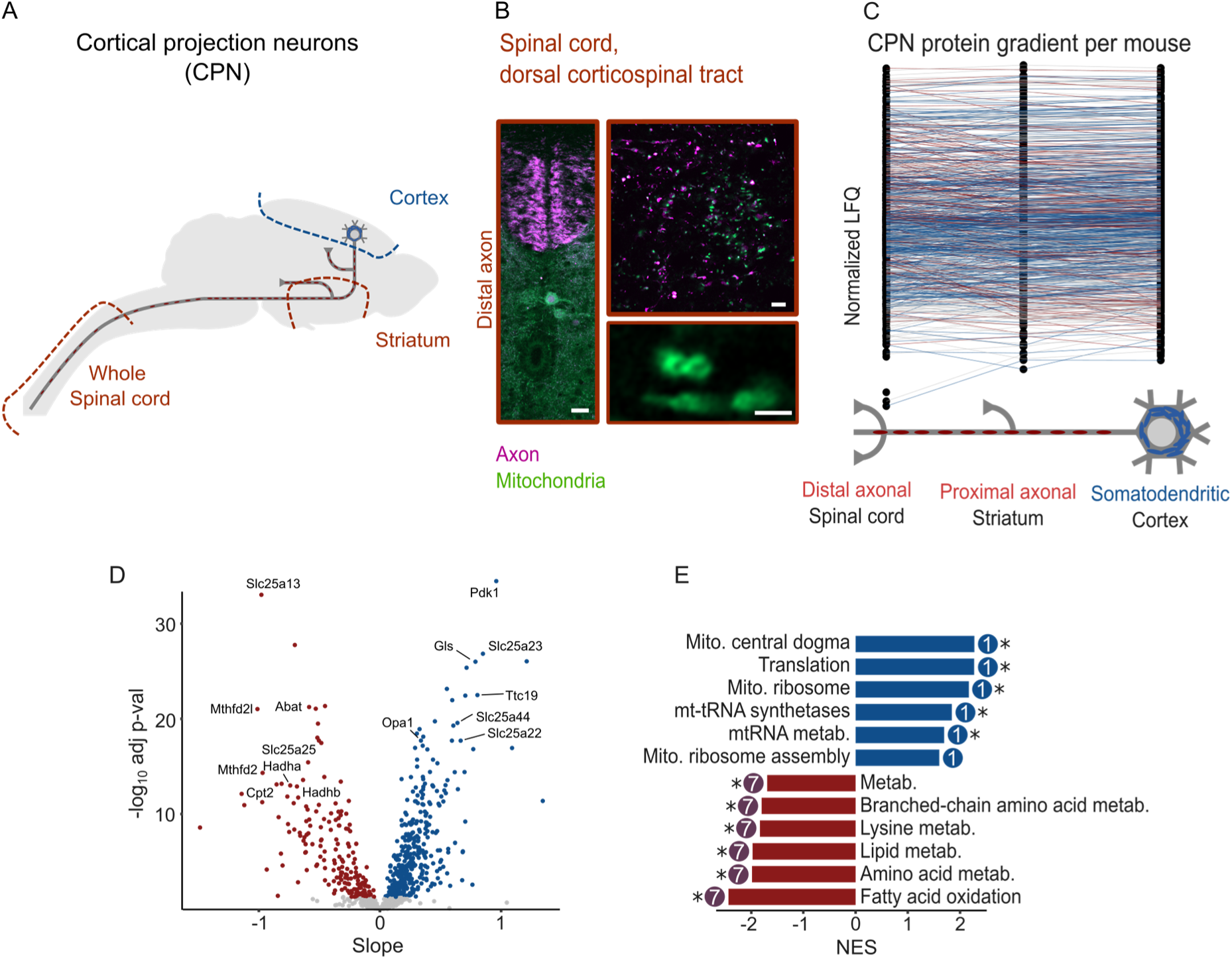
Protein gradient analysis. **A.** Mitochondria from CPN neurons were also collected from distal axons in the spinal cord. **B.** Airyscan transversal images of the dorsal corticospinal tract in the spinal cord of CPN; Rbp4-Cre driven GFP-OMM (green) and tdTomato (magenta). Scale bars: left overview panels 25 µm; right panels 5 µm (upper zoom), 1 µm (lower zoom detail). **C.** Normalized mitochondrial protein abundances, colored by slope direction as per panel D: positive slope (higher in soma, blue), negative slope (higher in distal axon, red), or flat (unchanged, gray). **D.** Linear regression analysis (volcano plot) of protein gradients shown in panel C; significance defined as FDR < 0.05. **E.** Gene set enrichment analysis using MitoPathways3.0. Somatodendritic pathways are shown in blue; axonal pathways in red. Pathways are labeled with their group from Fig. 2H. * indicates FDR < 0.05. Somatodendritic and proximal axon data also shown in Fig. 2.

**Supp. Fig. 4:**
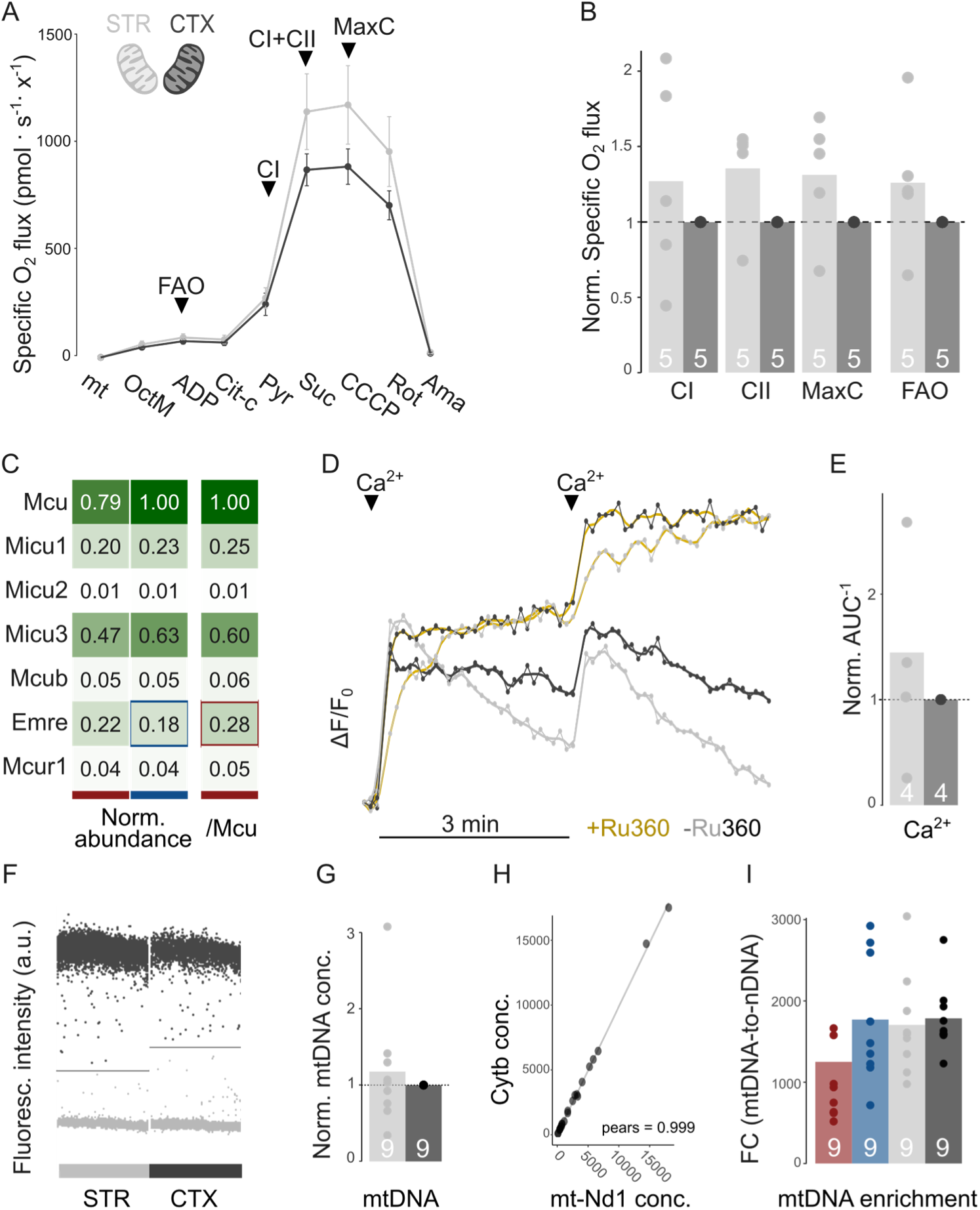
Complementary data to functional validation and mtDNA quantification. **A.** Respirometry curves of mitochondria isolated from cortex (CTX; background for CPN somatodendritic compartment; black) and striatum (STR; background for CPN axonal compartment; grey). Data are shown as means ± SEM. **B.** Pairwise normalized replicates from panel A. CI: complex I; CII: complex II; MaxC: maximum capacity; FAO: fatty acid oxidation. **C.** Proteomic log_2_ fold changes of the Mcu complex in the CPN dataset. **D.** Representative mitochondrial calcium uptake curve in the absence (black: cortex; grey: striatum) and presence of the Mcu inhibitor Ru360 (gold). Arrowheads indicate calcium addition. **E.** Pairwise normalized replicates of the calcium uptake assay. **F.** Representative digital PCR quantification of mitochondrial DNA (mtDNA), showing separation of positive and negative partitions in cortical and striatal mitochondria. The fluorescence threshold indicated by a horizontal line separates positive (black) and negative (gray) partitions. **G.** Pairwise normalized replicates from mtDNA quantification assay. **H.** Correlation of mtDNA quantification with mt-Nd1 and Cytb loci. **I.** Fold-change (FC) enrichment of mtDNA (probed with mt-Nd1 primers) relative to nuclear DNA (nDNA; probed with Actb primers) in isolated mitochondria. Related to panel G and Fig. 3F. Norm.: normalized; Conc.: concentration. Color coding: blue, somatodendritic mitochondria from CPN; red, axonal mitochondria from CPN; black, cortical (CTX) mitochondria; grey, striatal (STR) mitochondria.

**Supp. Fig. 5:**
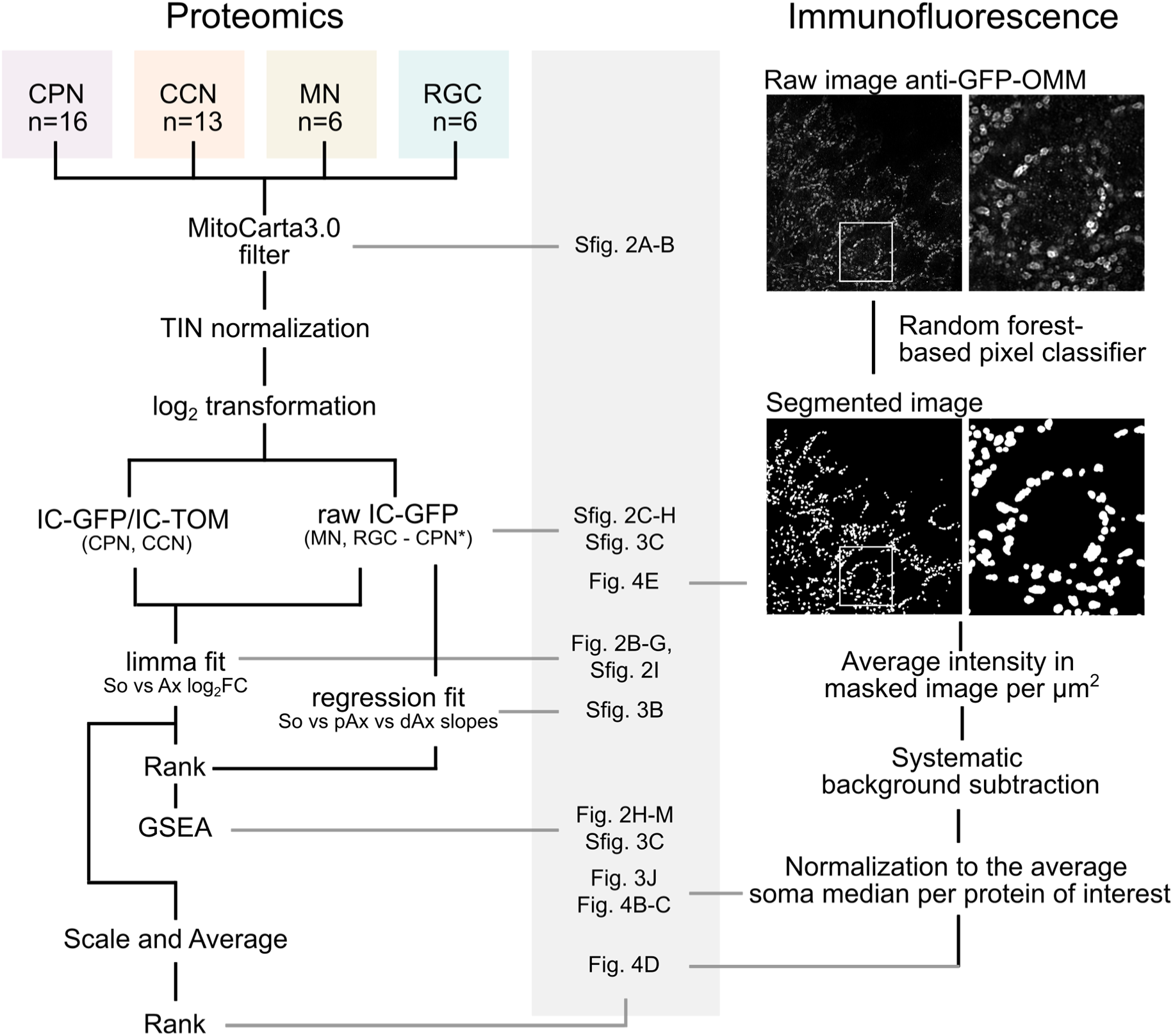
Schematic explanation of data analysis. Step-by-step proteomics and immunofluorescence data processing explaining at which data-processing step figure panels were produced. CPN: cortical projection neurons. CCN: central cholinergic neurons. MN: motor neurons. RGC: retinal ganglion cells. TIN: Total intensity normalization. IC-GFP/IC-TOM: Immunocapture with GFP or TOMTOM22 antibody beads. So: Soma. Ax: Axon. pAx: proximal axon. dAx: distal axon. GSEA: Gene Set Enrichment Analysis. *For protein gradient analysis.

